# Monocytes are biological sensors of aging and frailty in humans

**DOI:** 10.64898/2026.03.10.710601

**Authors:** Chanhong Min, Charles Ezenwanne, Yoseph W. Dance, Nico Macaluso, Ladaisha Thompson, Lolita Nidadavolu, Akhil Katuri, Crystal Szczesny, Jacqueline Langdon, Edward J. Pearce, Peter Abadir, Jeremy D. Walston, Jude M. Phillip

## Abstract

Among older adults, frailty is clinically identifiable and characterized by increased vulnerability to adverse health outcomes. Although various tools exist to identify frailty, diagnosis often depends on late-stage clinical symptoms, which typically limit proactive intervention options. We considered whether aging and frailty information may be encoded in the single-cell behaviors and dynamic responses of primary human monocytes. Combining high-content imaging, single-cell behavior profiling, and machine learning, we demonstrated unique age- and frailty-dependent monocyte behaviors at baseline and following exposure to inflammatory stressors. Using these single-cell behaviors, we developed a deep learning neural network model called scTRAIT. scTRAIT accurately predicts the frailty status of older donors, including the capability to track and forecast longitudinal changes in frailty status. Collectively, these findings demonstrate that aging and frailty information are robustly encoded within single-cell behaviors, establishing monocytes as significant biological sensors of aging and frailty in humans.

**Teaser:** Single-cell monocyte responses enable robust prediction of aging and frailty in humans

## INTRODUCTION

Many older adults endure multiple chronic conditions daily, facing elevated risks of physical and cognitive decline^1,2^. Yet the health status and intrinsic capacity for coping with such conditions vary widely among older adults. Understanding the scope of this variability is vital given the ongoing shifts in global population demographics towards an increasing fraction of adults aged 60 and older^3,4^. This shift is poised to considerably impact the socioeconomics of multiple countries by creating a disparity between maintaining quality of life amidst rising healthcare costs^5–7^. Older adults are at the greatest risk for developing multiple chronic diseases in a progression towards poor health^8^. As such, there is an urgent need to develop novel approaches to monitor health, predict and stratify which older adults may be most vulnerable to the adverse effects of aging, and identify those who could benefit most from targeted interventions.

A major challenge rests in the reactive nature of our current medical system, which often prioritizes the management of age-related dysfunctions rather than anticipating them for targeted intervention and prevention^9^. Key to this situation is the current lack of cheap and effective tools to accurately measure baseline health, predict individual aging trajectories, and ultimately forecast adverse aging and the emergence of age-related conditions such as frailty. Frailty is clinically identifiable and meaningful, characterized by reduced physiological reserve and an increased vulnerability to physiological stressors such as an injury or a fall^10,11^. This increased vulnerability can lead to severe health consequences, such as susceptibility to illness, prolonged hospital stays, and death^12–14^. In a recent study of community-dwelling older adults across sixty-two countries, 11% of adults aged 50-59 were frail, which rose to 51% among adults aged 90 and older, highlighting the growing prevalence of frailty with increasing age^15^.

Although several methods exist to identify and diagnose frailty among older adults, two clinical classification concepts dominate: the physical frailty phenotype and the frailty index^10^. The physical frailty phenotype defines frailty as a syndrome characterized by a collection of symptoms, including weakness, slowness, physical inactivity, exhaustion, and unintentional weight loss. In this paradigm, a person is considered robust or non-frail if they exhibit none of these frailty characteristics, pre-frail if they present with 1 or 2 characteristics, and frail if they present with 3 to 5 characteristics. Further, a person presenting with all 5 frailty characteristics is considered severely frail^16^. A study evaluating the prevalence of frailty among older adults, using the physical frailty phenotype, found that in the United States, ∼15% of older adults aged 65 and older were considered frail, with another ∼45% being considered prefrail^17^.

Alternatively, the frailty index defines frailty as a state of poor health characterized by the accumulation of health deficits^18,19^, which are broadly defined based on health surveys, geriatric assessments, and/or molecular biomarkers^20^. By definition, the frailty index ranges from 0 to 1. A person with a frailty index of 0 exhibits no measured deficits and, hence, is considered non-frail. Persons with a frailty index of 0.7 or higher, meaning that they have >70% of the measured health deficits, are considered severely frail^10^.

While both methods identify frailty based on fundamentally different metrics that do not always correlate, they both rely on physiologically observable, late-stage symptoms^14,21^. This reliance on late-stage indicators, such as poor gait time and grip strength, accumulation of health deficits, or chronic low-grade inflammation, presents limitations for proactive clinical decision-making and identifying timely interventions to slow or prevent frailty progression for improving health^22,23^. This is further exacerbated by the qualitative and static nature of most of the defining criteria, providing limited insights into an individual’s physiological reserve or dynamic capacity. We considered whether more quantitative and dynamic cellular measures could provide a solution, especially since dysfunctions arise in cells well before they are clinically apparent.

Previously, we have shown that aging information is encoded within the biophysical properties of cells, controlling, for example, their ability to move, change their morphology, and exert forces as they navigate complex microenvironments^24^. Even when cells are taken out of their natural physiological context, they retain a memory of where they originated and can modulate their responses and/or behaviors based on the health and age of the individual from which they were sampled^25,26^. However, not all cells exhibit the same predictive potential to inform health, aging, or the emergence of disease.

Immune cells undergo profound changes during healthy aging and frailty, driven in part by molecular and cellular compositional changes that give rise to immunosenescence and chronic low-grade inflammation (inflammaging)^23,27^. Monocytes, a heterogeneous population of innate immune cells, are central mediators of inflammation, playing key roles in the development of frailty-associated phenotypes ^28,29^. With age, monocytes exhibit substantial phenotypic and functional alterations, including shifts in subset composition, adhesion, chemotaxis, phagocytosis, pro-inflammatory cytokine production, receptor signaling, and differentiation patterns, which collectively shape how they dynamically behave and respond to perturbations ^30–33^. Here, we hypothesize that monocyte behaviors and the magnitude of their responses to defined perturbations and stressors could provide biologically meaningful readouts to proactively determine health, aging, and frailty in humans.

## RESULTS

### Single-cell analysis reveals distinct age-associated patterns of motility among young, old, and frail monocytes

Cells encode information related to the health and aging of individuals, which can be measured quantitatively based on biophysical properties that describe how cells move, change their morphologies, and exert forces as they navigate complex microenvironments^24,25^. To determine whether primary monocytes encode aging and frailty information, we recruited a balanced cohort of forty-five donors through an established donor registry within the Johns Hopkins Older Americans Independence Center. Our cohort comprised three groups: young adults (aged 18-47), non-frail older adults (aged 65-90), and frail older adults (aged 65-90). Among the older adults, we determined frailty status using the physical frailty phenotype criteria^16,22^. From each donor, we drew whole blood, purified peripheral blood mononuclear cells, isolated primary CD14^+^ monocytes, and performed single-cell motility profiling and analysis (**Figure 1A**, see Methods). Throughout this report, we refer to monocytes from young donors as young monocytes, monocytes from non-frail older adults as old monocytes, and monocytes from frail older adults as frail monocytes.

**Figure 1.**
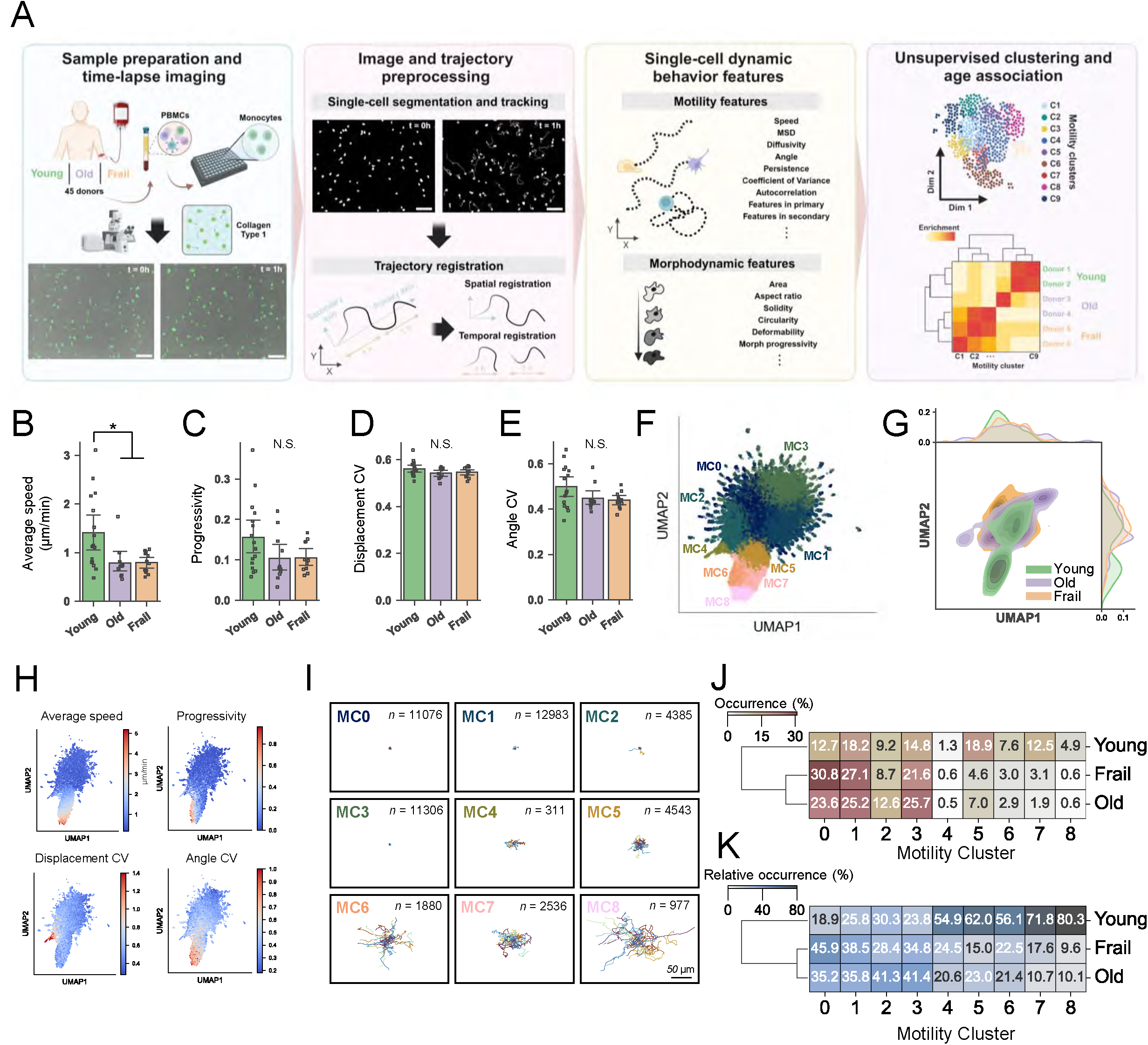
Baseline single-cell monocyte motility encodes age information. **A**. Schematic illustration for profiling single-cell monocyte behaviors. Scale bar denotes 80 µm. We performed live-cell imaging on monocytes from 45 donors, followed by single-cell segmentation, tracking, and trajectory registration. Approximately 100 motility and 30 morphodynamic features were extracted to characterize behaviors. Unsupervised clustering identified distinct motility states linked to age. **B-E**. Bulk analysis for average speed (B), progressivity (C), displacement coefficient of variance (D), and turning angle coefficient of variance (E) across age groups (mean ± 95% C.I.). Statistics analyzed via Kruskal-Wallis test with Dunn’s post-hoc test (young N=17 donors, old N=13 donors, frail N=13 donors;*p<0.05, N.S. means non-significant). **F**. 2D UMAP representation of nine motility clusters identified by unsupervised k-means clustering. **G**. 2D UMAP kernel density estimation (KDE) displays the density of single-cell per age-group across the manifold. **H**. UMAP feature plot denoting the single-cell magnitudes of average speed, progressivity, displacement CV, and angle CV. **I**. Thirty randomly selected movement trajectories per MC, total cells in each MC are indicated. **J**. Heatmap showing the fractional occurrence of nine MCs across the three age groups. Hierarchical clustering is based on the Euclidean distance using the Ward method (young n=3,169 cells, old n=3,597 cells, frail n=5,508 cells). The sum of rows equal 1. **K**. Heatmap showing the fractional occurrence normalized by each MC across age groups. Hierarchical clustering is based on the Euclidean distance using the Ward method (young n=3,169 cells, old n=3,597 cells, frail n=5,508 cells). The sum of columns equals 1.

To determine whether young monocytes exhibit distinct motility characteristics relative to old or frail monocytes, we quantified bulk motility based on the average of each motility feature (e.g., total distance travelled and monocyte speed) per donor across the young, old, and frail groups. We observed that at baseline, young monocytes exhibited enhanced motility relative to old and frail monocytes (**Supplementary Figure 1A**). For example, young monocytes travelled ∼60% faster than old or frail monocytes, which was consistent with previous findings of decreased bulk cell motility with increasing age (**Figure 1B**)^34,35^. However, among the three groups, there was no significant difference in the progressivity that measures the directedness of a cell’s movement (**Figure 1C**), and both the displacement and angle coefficient of variation (CV) (**Figure 1D-E**). Notably, the bulk motility features did not distinguish clear differences between old and frail monocytes.

Because monocytes are inherently heterogeneous, we considered whether single-cell analysis could provide additional information that was not directly apparent from the bulk analyses. To investigate this, we conducted single-cell motility analysis using the custom computational workflow, designated CaMI, developed in our laboratory^36^. CaMI is a machine-learning framework that captures the heterogeneous motility behaviors of single cells and treats each cell as a single entity to find groupings of cells based on similarities and differences across multidimensional motility features. We refer to these groupings of single cells as motility clusters (MCs). To conduct the CaMI analysis, we pooled all individually tracked monocyte trajectories across all donors into a single database and conducted clustering analysis (see Methods). The results showed the emergence of nine distinct motility clusters. For ease of interpretation, we ordered the MCs based on the magnitude of the total displacements per cluster and visualized them in a two-dimensional space (**Figure 1F**). Detailed characterization of each MC highlighted distinct differences in the magnitudes of motility features (**Supplementary Fig. 1B**).

To visually interpret the motility space and estimate the spatial abundance of monocytes across the MCs, we projected the locations of young, old, and frail monocytes as density contours onto the motility space. While most of the old and frail monocytes occupied the upper portions of the UMAP, described by MC0-MC4, a large portion of young monocytes occupied the lower portion of the UMAP, described by MC5-MC8 (**Figure 1G**). Notably, monocytes in the lower portion of the space exhibited enhanced motility, with faster and more directed movements, as indicated by higher average speeds and progressivity, respectively (**Figure 1H**). These cells also exhibited low coefficients of variance in displacement and high coefficients of variance in the turning angles (**Figure 1H, Supplementary Figure 1C**). To further confirm this, we randomly selected thirty movement trajectories per MC and replotted them with all centered initially at the origin (**Figure 1I**). Qualitatively, we observed that MC0-MC4 described monocytes with little to no motility, while MC5-MC8 described monocytes with significantly higher motility. MC6 and MC8 were the most directed in their movements with high progressivity.

Quantifying the fractional abundance of monocytes from each of the three groups (young, old, and frail) across each MC, we again observed strong age associations and differences in cellular heterogeneity per group. Notably, young monocytes were most heterogeneous, while old and frail monocytes showed similarly lower levels of heterogeneity, as measured by the Shannon entropy (**Supplementary Figure 1D**). Furthermore, under 15% of old and frail monocytes (12.4% and 11.2%, respectively) were characterized by the high motility clusters MC5-MC8, compared to almost 45% of young monocytes being characterized in these high motility clusters (**Figure 1J**). To assess similarity in MC distributions across age groups, we computed pairwise cross correlations and found that young monocytes diverged from old (r = 0.62) and frail (r = 0.57), whereas old and frail monocytes were highly correlated (r = 0.96) (**Supplementary Figure 1E**). Additionally, based on the composition per MC, over 50% of monocytes in MC4-MC8 were in the young category (**Figure 1K**).

Comprehensively, the single-cell analyses revealed heterogeneous, age-associated motility patterns described by distinct motility clusters. However, even with the insights drawn from the cellular heterogeneity analyses, the single-cell motility patterns at baseline were unable to discern significant differences between old and frail monocytes.

### Dynamic responses to pro-inflammatory perturbations can distinguish young, old, and frail monocytes

Since frailty is associated with chronic low-grade inflammation, where cells are constantly exposed to inflammatory factors, we hypothesized that frail monocytes would exhibit reduced overall responses to pro-inflammatory molecules, relative to young or old monocytes. To assess this, we exposed the monocytes to 1 µg/mL of cell-free poly(dA-dT) double-stranded DNA, 100 ng/mL of recombinant human interleukin-6 (IL6), and 10 µg/mL of lipopolysaccharide (LPS) from *Escherichia coli*, then performed high-content live-cell imaging and analysis (**Figure 1A**).

We found that young monocytes displayed increases in cell motility features, such as average speed and progressivity, when exposed to cell-free DNA and IL6. In contrast, LPS induced a significant decrease (∼50%) in both the average speed and progressivity in young monocytes (**Figure 2A**). In line with the findings in **Figure 1**, old and frail monocytes exhibited reduced average speeds relative to young monocytes, as observed in control conditions. Old monocytes exhibited little to no responses when exposed to cell-free DNA and LPS but displayed enhanced motility to IL6 (Cohen’s D value greater than 0.5) (**Figure 2B**). In line with our initial hypothesis, frail monocytes exhibited no response to any of the inflammatory perturbations, suggesting an inherent insensitivity to inflammatory cues and a potential decoupling between the sensing of inflammatory perturbations and modulating their motility (**Figure 2C**). To expand the insights beyond the speed and progressivity, we calculated the log fold-change for the expanded set of motility parameters relative to the control for each group, then curated the motility features with changes greater than 30% (**Figure 2D**). This expanded list of cell motility features further supported the findings and suggests that while young and old monocytes exhibit high or moderate responses to inflammatory perturbations, frail monocytes are characterized by distinct insensitivity to the panel of inflammatory perturbations evaluated.

**Figure 2.**
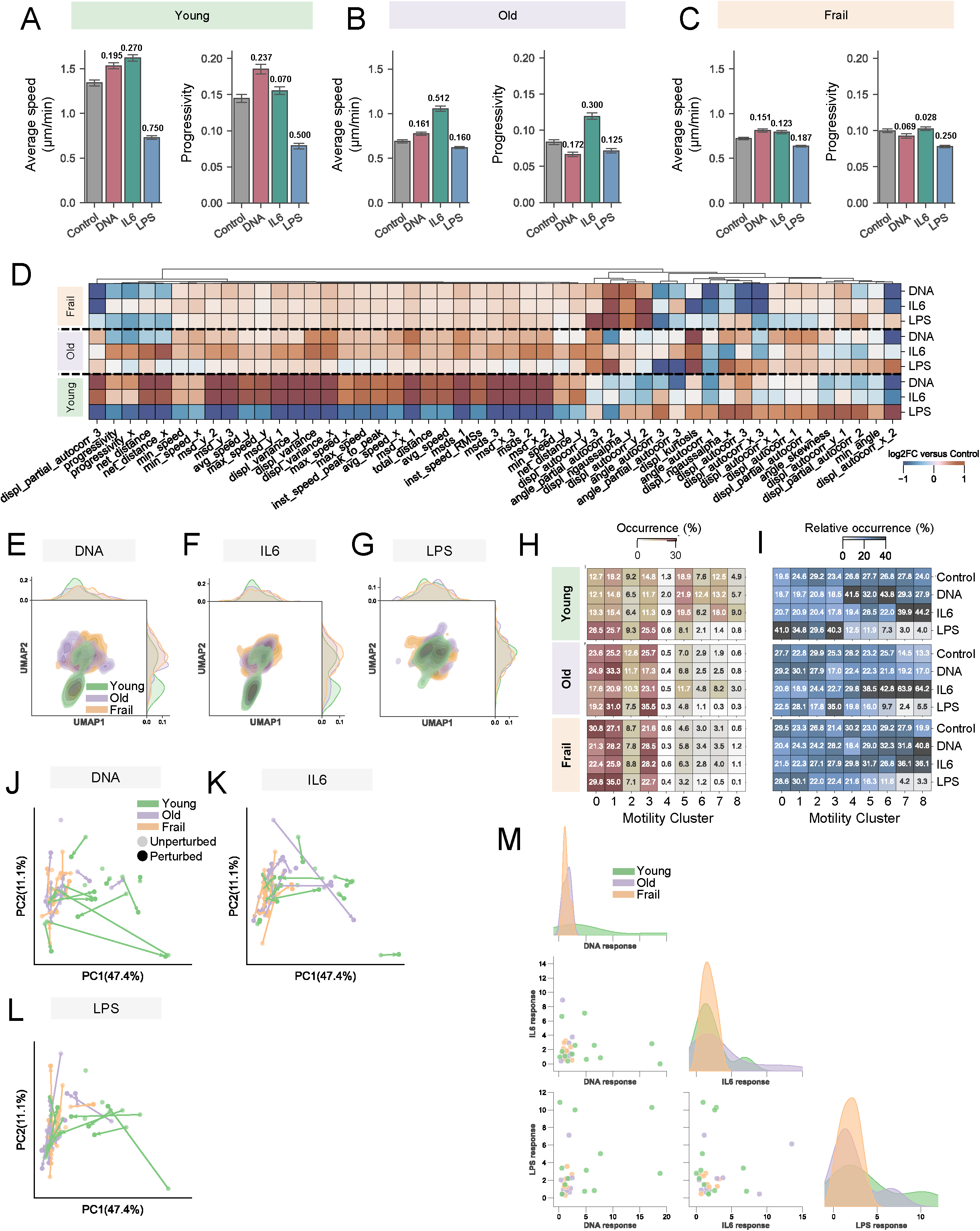
Monocytes respond to inflammatory stressors in an age- and frailty-dependent manner. **A-C**. Left, total distance traveled per hour (µm) and right, progressivity computed across vehicle control and three perturbation conditions. Young monocytes (A) exposed to: Control n=3,169 cells, DNA n=2,795 cells, IL6 n=3,362 cells, and LPS n=2,370 cells. Old non-frail monocytes (B) exposed to: Control n=3,597 cells, DNA n=3,454 cells, IL6 n=3,422 cells, LPS n=3924 cells. Frail monocytes (C) exposed to: Control n=5,508, DNA n=4,949 cells, IL6 n=5,399 cells, LPS n=6,219 cells. Annotations show absolute Cohen’s d values (small effect size, d≤0.2; large effect size, d≥0.5), each compared to the control. Error bars denote the mean ± 95% confidence intervals. **D**. Heat map showing the relative log fold changes over control for select motility parameters (bottom) express more than 30% change over the control in at least one condition. Dendrogram on top denoted unsupervised hierarchical clustering of parameters based on average correlation distances. **E-G**. 2D UMAP KDE representation of motility space across three age groups exposed to DNA (E), IL6 (F), and LPS (G) conditions. 1D distributions of single cells for each at the top and right margins. **H**. Heatmap of fractional occurrence of nine MCs across the four conditions for each age group. **I**. Heatmap of fractional occurrence normalized by each MC across four conditions for each age group. **J-L**. Principal component analysis (PCA) plot showing the responses of 16 donors in young, 13 donors in old, 12 donors in frail (41 donors in total) in the absence or presence of perturbation conditions. Each connected vector depicts the average trajectory relative to control for monocytes exposed to DNA (J), IL6 (K), LPS (L). **M**. Magnitude of perturbation response plot across all donors in the age group. Off-diagonal panels indicate pairwise magnitude of perturbation response quantified across all donors. Diagonal panels show KDE distribution of single condition magnitudes of perturbation responses.

To effectively profile the heterogeneity of responses, we again conducted single-cell analysis of monocytes exposed to cell-free DNA, IL6, and LPS, then projected them onto the previously defined motility space (**Figure 1F**). Monocytes exposed to DNA and IL6 displayed distinct, age-related shifts in their motility (**Figure 2E-G**). Young monocytes exposed to DNA and IL6 induced heterogeneous motility patterns characterized by shifts towards the high motility regions (low UMAP2) (**Figure 2E-F**). Intriguingly, LPS exposure across all conditions led to a convergence of motility patterns, which were primarily described by little to no motility (center of UMAP) (**Figure 2G**). This convergence of motility in response to LPS suggests that this potent inflammatory signal could override key age-related monocyte motility and induce a uniform inhibition of cell movements. Computing the abundance of cells within each MC more clearly demonstrated the effect of LPS exposure, displaying an increased abundance in low motility clusters (MC0, MC1, and MC3) in the young monocytes. IL6 treatment of old monocytes resulted in a modest increase in the abundance of cells in MC5 and MC7 (**Figure 2H**). Furthermore, over 40% of old monocytes in MC6–MC8 were from the IL6-treated group. This was consistent with the prior observation that IL6 induced enhanced cell motility and sensitivity (**Figure 2B**). In contrast, frail monocytes showed no clear dominance of any condition, further supporting their insensitivity (**Figure 2I**).

Inter-person differences may have contributed to the variations we found between age groups. We observed heterogeneous MC distributions and hierarchical clustering across donors with varying ages (**Supplementary Figure 2A–C**). Young monocytes generally exhibited large response vectors along the high-variance principal component 1 (PC1) following stimulation, whereas frail monocytes displayed attenuated PC1 responses (**Figure 2J–L**). This was further demonstrated by higher perturbation response in young monocytes and narrow, insensitive responses in frail monocytes across donors (**Figure 2M**). To exclude sampling bias as a confounding factor of age- and frailty-associated patterns, reconstruction of the motility space using equal numbers of monocytes per donor recapitulated the original space geometry (**Supplementary Figure 3A**), preserved age-group clustering and perturbation-specific shifts, including IL6-associated increases in MC5 and MC7 in old monocytes (**Supplementary Figures 3B-I**), and showed high concordance with the original space (**Supplementary Figures 3J-M**). Another confounding covariate, biological sex, exerted minimal effects in control conditions but significantly modulated motility in old and frail monocytes, with males exhibiting higher speed and progressivity than females (**Supplementary Figure 4A-B**) and sex-specific differences in MC heterogeneities, which were condition-dependent (**Supplementary Figure 4C-F**).

Collectively, these findings demonstrate that while young monocytes exhibit robust responses to inflammatory perturbations, old monocytes show sensitivity to IL6, and frail monocytes show insensitivity across all the inflammatory perturbations evaluated. These results highlight an inherent relationship between single-cell monocyte responses and a person’s age and frailty status.

### Age- and frailty-related monocyte motility depends on the local cell density

Monocytes are known to adapt their motility in response to local microenvironmental cues. For example, at high densities, monocytes exhibit increased movements partly due to the increased secretion of IL6^37^. This finding prompted us to investigate whether specific attributes of local monocyte communities influence the age- and frailty-related monocyte motility. Although the same number of cells were seeded, when monocytes moved, they formed local cell communities with varying population densities. We analyzed these spatial localization patterns from two perspectives (**Figure 3A**). First, we quantified the local cell density by counting the number of neighboring cells that each monocyte encountered within a 100 μm search radius to assess how local cell density influenced single-cell motility (**Figure 3A**, left). Second, we quantified cell-cell distances by constructing a spatiotemporal distance tensor to determine nearest-neighbor distances (**Figure 3A**, right). Overlaying local density maps onto tracked monocyte trajectories revealed qualitative spatiotemporal variations and patterns of monocyte motility as a function of local monocyte density (**Figure 3B**).

**Figure 3.**
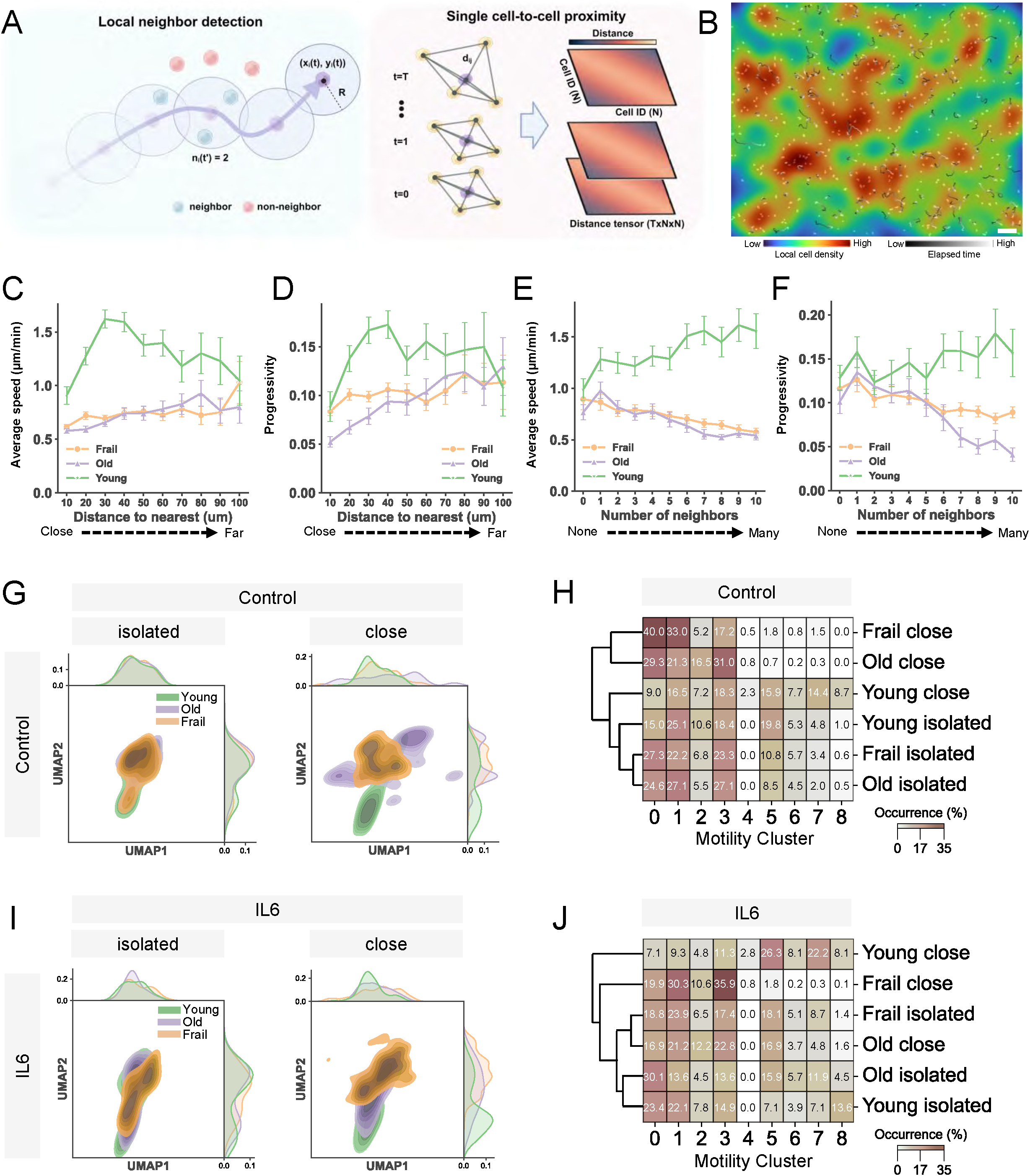
Cell-to-cell spatial proximity exhibits divergent monocyte motility based on age and frailty status. **A**. Schematic illustration of spatial colocalization analysis based on two algorithms: Left, local neighbor detection assigns neighbor cells following the cell movement trajectory. n(t), number of neighbor cells at given time t; R, search range radius; (x(t), y(t)), location of the moving cell at time t. Right, the distance tensor is constructed based on the cell-to-cell pairwise distances using the time evolution of cell spatial centroids. **B**. Color-coded image of single-cell movement trajectories, trails denote the distance traveled per cell, and the colors indicate the local cell density map. Only trajectories tracked for more than 20 frames are shown. Scale bar: 100 µm. **C-D**. Line plots showing how the average speed (G) and progressivity (H) changes based on the distance to the nearest cell with 10 µm bin (mean ± 95% C.I.). **E-F**. Line plots showing how the average speed (I) and progressivity (J) changes based on the average number of neighbors within 100 µm per cell. Error bars denote mean ± 95% confidence intervals. **G**. 2D UMAP KDE representation for isolated cells and cells in close proximity within the control condition. Distributions of single cell abundances are denoted at the top and right margins. Left, isolated cells (average neighbor ≤ 0.6 cells); right, close cells (average neighbor ≥ 9 cells). **H**. Heat map showing the fractional occurrences of cells per MCs across age groups for isolated and close conditions. Dendrogram shows hierarchical clustering of Euclidean distances with Ward linkages (Young isolated n=207 cells, Young close n=611 cells, Old isolated n=199 cells, Old close n=1,154 cells, Frail isolated n=176 cells, Frail close n=1,029 cells). **I**. 2D UMAP KDE representation for isolated cells and cells in close proximity within the IL6 condition. Distributions of single cell abundances are denoted at the top and right margins. Left, isolated cells (average neighbor ≤ 0.6 cells); right, cells with many neighbors (average neighbor ≥ 9 cells). Dendrogram shows hierarchical clustering of Euclidean distance with Ward linkages (Young isolated n=154, Young close n=505 cells, Old isolated n=176 cells, Old close n=189 cells, Frail isolated n=138 cells, Frail close n=1,210 cells).

Next, we quantified the age and frailty-associated effects of local cell density on monocyte motility. For young monocytes, we observed a biphasic relationship between the nearest cell neighbor distance and the monocyte’s average cell speed and progressivity, with peak values when cells were within approximately 30-40 μm of neighboring cells. Conversely, old and frail monocytes exhibited a gradual increase in their average speed and progressivity, with similar magnitudes observed for young, old, and frail monocytes when they were far away (approximately 100 μm) from the nearest cell (**Figure 3C, D**). Similarly, young monocytes exhibited a strong increase in average speed as the number of cell neighbors increased, likely from increased protein secretions (e.g., IL6) and other cell-cell interactions. Interestingly, old and frail monocytes decreased their average speed with an increasing abundance of neighbors, suggesting that a high-density context was inhibitory to their movements (**Figure 3E**). This trend was mirrored for progressivity, but with a high number of neighbors (between 7-10 cells), frail monocytes exhibited less progressive (more tortuous) movements than the old (**Figure 3F**). These were observed at baseline and with exposure to pro-inflammatory perturbations, with the only exception being LPS (**Supplementary Figure 5A-C**).

To further investigate the effects of spatial localization effects, we categorized cells into two groups: “close”, which we defined as monocytes with nine or more neighbors detected, and “isolated”, defined as those encountering up to one neighbor within the 100 μm search range for the tracked duration. Remarkably, at baseline, isolated monocytes displayed a convergence in motility cluster distributions across all age groups, indicating a similar motility profile when isolated, irrespective of age or frailty (**Figure 3G**, left). This suggests that when monocytes are spatially isolated (i.e., far away from other monocytes), their motility becomes less indicative of age or frailty status, possibly due to reduced interactions with neighboring cells. In contrast, “close” monocytes exhibited differential shifts in the patterns of MCs based on the local cell density (**Figure 3G**, right). Hierarchical clustering of age- and frailty-related MC distributions further confirmed these observations, showing that “isolated” monocytes maintained similar MC distributions across age groups. However, “close” monocytes demonstrated significant shifts in their motility across age groups, characterized by a higher fraction of monocytes in MC5-MC8 for young, MC0-MC3 for old, and MC0, MC1, and MC3 for frail (**Figure 3H**). These results indicate age- and frailty-specific sensitivities at high local cell densities, which are likely due to increased cellular interactions and local environmental cues, such as secretions and physical cell-cell contacts, which influence monocyte functions and inflammatory responses.

In light of these cell-density influences, we investigated whether exposure to pro-inflammatory perturbations also modulates the impact of local cell density on age- and frailty-related cell motility. Notably, IL6 exposure showed a high overlap in the motility distributions of “isolated” cells across age groups. However, monocytes across all age groups maintained distinct motility when in “close” context (**Figure 3I**). These IL6-induced shifts in the motility (**Figure 3I**, right) were different from our observations of monocytes in “close” conditions at the baseline/control context (**Figure 3G**, right). Hierarchical clustering of the MC distributions across young, old, and frail monocytes confirmed these results and further indicated that in “close” context, young and frail monocytes are most divergent. For example, while young monocytes exhibited more diverse motility patterns, frail monocytes converged towards low motility MCs (MC0-MC3) (**Figure 3J**). Similarly, DNA and LPS exposures also influence the effects of local cell density on monocyte motility (**Supplementary Figure 6A, B**). Notably, cross-correlation analysis based on MC distributions revealed that young monocytes in “close” conditions were most different from all other conditions, which held true across all exposures (**Supplementary Figure 6C-F)**.

Collectively, these findings reveal that local cell density significantly influences monocyte motility. Furthermore, pro-inflammatory perturbations can modulate the effects of local cell density, highlighting that local cell density and inflammatory conditions jointly modulate the single-cell motility of monocytes.

### Age- and frailty-dependent features define the relationships between monocyte behaviors and chronological age

We investigated whether cell behaviors reflected the chronological age of donors by quantifying their correlations with age and frailty status. Single-cell behaviors refer to the biophysical responses of monocytes to inflammatory perturbations, primarily describing multiple characteristics such as cell motility and changes in cell morphologies (morphodynamics). Initial analyses showed that the chronological ages of young donors were positively correlated with average speed and progressivity, while old donors exhibited negative correlations for both baseline and IL6-exposed young monocytes. However, frail donors displayed weak correlations, suggesting a decoupling of cell motility and chronological aging in frail donors (**Supplementary Figure 7A, B**).

Beyond average speed and progressivity, we evaluated age correlations with behavioral attributes consisting of multiple motility and morphodynamic features. The analysis revealed distinct age- and frailty-related patterns (**Figure 4A**). Young donors displayed a strong positive correlation between behavioral features and chronological age, except for DNA perturbation. In contrast, monocytes from old donors showed a significant negative correlation between behavioral features and chronological age, which persisted under control conditions and all three pro-inflammatory perturbations. However, frail donors displayed weak to negligible correlations across all conditions.

**Figure 4.**
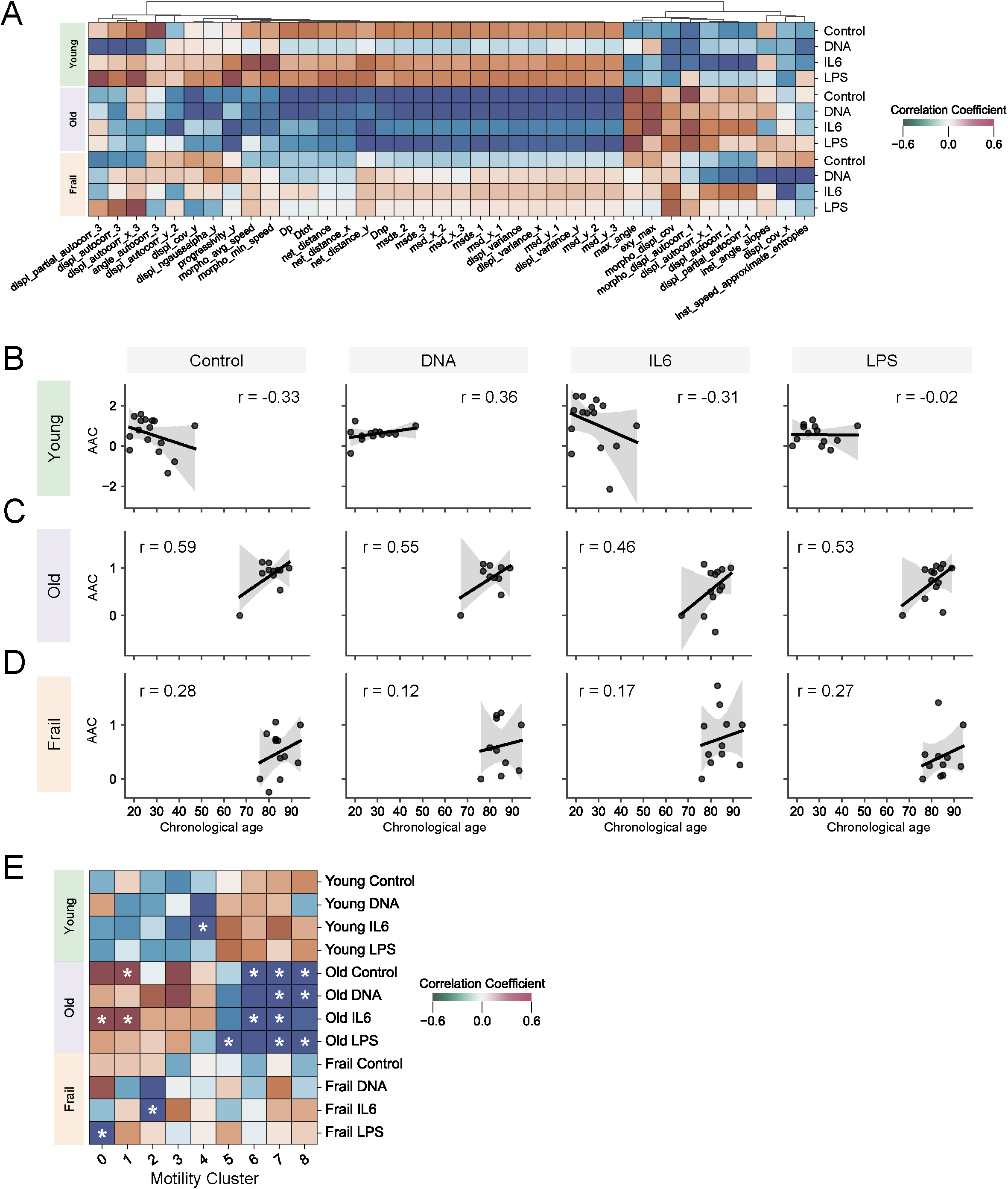
Behavioral features reveal distinct age- and frailty-dependent correlations with chronological age. **A**. Heatmap showing the Pearson correlation coefficients of select behavior parameters with chronological age. Behavior parameters were selected if at least one condition had a p-value less than 0.05 in any age group. The dendrogram denotes hierarchical clustering of parameters based on the average correlation distances. **B-D**. Piece-wise regression plots per age-group showing the relationship between the age axis coefficient and chronological age across perturbation conditions; Young (B), Old (C), and Frail (D). r indicates Pearson’s correlation coefficient. **E**. Heatmap showing the Pearson’s correlation coefficient between MC occurrence and chronological age across age groups and perturbation conditions. NOTE: * denotes conditions with p-values < 0.05.

In addition to assessing how single-behavior features encoded chronological age, we sought to develop an approach incorporating multi-dimensional motility behaviors that derive a comprehensive representation of cellular aging. To achieve this, we constructed a simple linear model that integrates multiple motility features into a single composite metric to capture relationships between cellular behaviors and chronological age^38^. Specifically, within each age group, we established a multi-dimensional linear age axis, with the youngest donor per age group serving as the initial point and the oldest donor per age group serving as the terminal point, thereby defining a linear trajectory of cellular motility behaviors across an age spectrum. We denoted this parameter as the age axis coefficient (AAC), representing the alignment of a donor’s cellular behaviors along the defined age axis (see Methods).

When applying this model, under both control and IL-6 conditions, we identified a negative correlation with the age axis among young donors, indicating a failure of the model (**Figure 4B**). This failure is likely driven by the high degree of variability we measured in cellular behaviors between young donors, suggesting a non-linear relationship between chronological age and cellular motility in this age group. This variability likely reflects the higher heterogeneity of cellular responses to perturbations occurring at younger ages, as evidenced by the positive correlation observed in the DNA results. Interestingly, DNA perturbation restored the correlation with the age axis, suggesting that specific stress conditions can correct for the otherwise unpredictable nature of motility behaviors in young donors. Conversely, LPS perturbation diminished the correlation, further indicating the context-specific nature of cellular motility in younger donors.

In contrast, with old donors, we observed consistently high positive correlations between the age axis coefficients and chronological age across all conditions (**Figure 4C**). This suggests that cellular motility features in old donors strongly encode chronological age, and these features maintain a stable relationship with age even when subjected to different inflammatory perturbations. This robustness in the monocyte motility-age relationship among the old donors reflects rigorous and predictable cellular behavior patterns, possibly due to the reduced heterogeneity associated with aging seen in our results. Frail donors presented a distinct profile, with correlations consistently weak across all perturbations (**Figure 4D**). This lack of significant correlation, despite the perturbations, suggests that frailty disrupts the encoding of age in monocyte behaviors, effectively decoupling cellular behavior from chronological aging. The weak correlations observed among frail donors suggest that frailty may represent a fundamentally different biological phenotype. Furthermore, this also suggests that frailty does not adhere to a typical linear aging trajectory, but rather that frailty reflects a distinct physiological state, which emerges independently of chronological age.

Extending our analysis to MC populations, we examined how the fractional abundance of specific MC correlated with chronological age (**Figure 4E**). Among old donors, we identified strong positive correlations between the fractional abundance of MC0 and MC1 and chronological age for control and IL6 conditions, indicating that MC0 and MC1 monocyte populations expand linearly with age. Conversely, we observed strong negative correlations between age and the fractional abundance of MC6, MC7, and MC8, suggesting a reduction in these subpopulations with increasing age. These consistent patterns were observed across both control and perturbed conditions, reinforcing the premise that cellular and population-level behaviors in old (non-frail) donors are tightly linked to chronological age. In contrast, frail donors exhibited few strong correlations with MC fractional abundances. Notably, MC0 and MC2 showed a degree of correlation under specific perturbation conditions, but overall, the strength of these correlations was significantly weaker than in the old donors. This result reinforces a disruption in normal aging processes associated with frailty and supports the hypothesis that frailty represents an age-independent phenotype.

### Deep neural network-based model predicts age and frailty status based on single-cell monocyte behaviors

Based on these findings, we considered whether conventional feature-engineering methods presented the best way to establish the relationships between monocyte behaviors, age, and frailty. To ensure a thorough analysis to describe single-cell behaviors, which accounts for features beyond those we could mathematically define, we implemented deep neural network (DNN) models. We chose DNNs because they can be effective in unraveling otherwise undiscernible, complex patterns within large datasets^39–44^. Thus, we built a DNN-based prediction model of age and frailty status. This model was also intended to validate findings identified in **Figures 1-4**, which showed that perturbation and colocalization-associated behaviors encoded aging and frailty status. To accomplish this, we developed **s**ingle-**c**ell **t**ime series and **r**esponse-**a**ided **i**nference **t**echnology—scTRAIT. scTRAIT couples selected single-cell behaviors with specific perturbation-responses to predict age and frailty at single-cell resolution. Conceptually, scTRAIT encodes the 3 behavior modalities (i.e., motility, colocalization or density-based effects, and morphodynamics) into respective latent vector representations. Each perturbation is then transformed into a multidimensional embedding that learns behavior-specific deviations induced by a particular perturbation. Once these embeddings are established, the model learns the post-perturbational behaviors of each modality and subsequently applies the correlations between the perturbed behaviors to predict age and frailty status (**Figure 5A**, see Methods). To benchmark the performance of scTRAIT, we prepared two additional baseline models to infer age and frailty status: A model based on movement trajectories (VT) and the other on morphodynamic features (VM) (see Methods).

**Figure 5.**
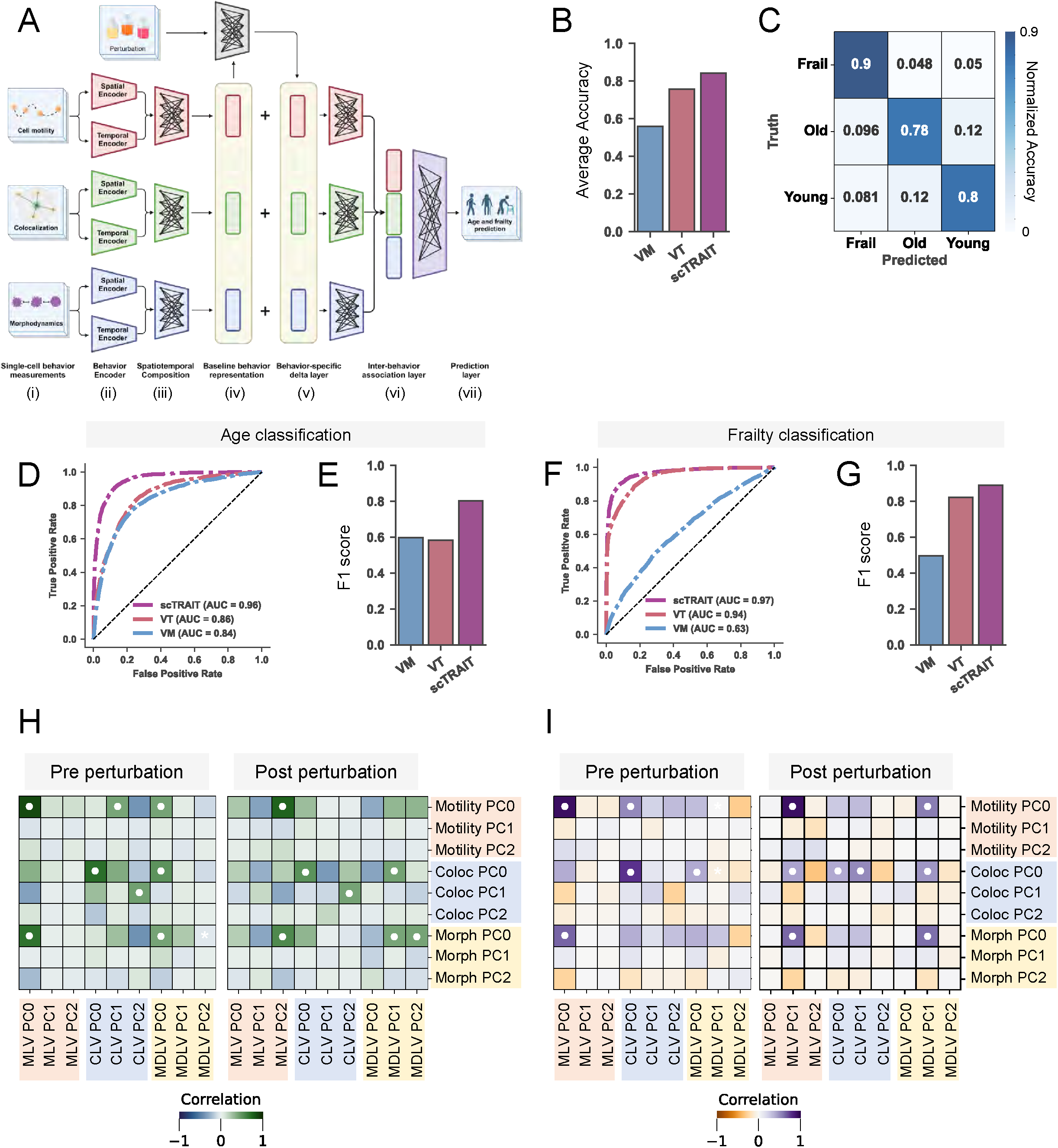
scTRAIT accurately predicts age group and frailty-status based on single-cell behaviors. **A**. Schematic of the scTRAIT model architecture for age and frailty prediction: (i) scTRAIT inputs movement trajectories, colocalization features, and morphodynamic descriptors per single cell. (ii) scTRAIT contains a spatial and temporal extractor, followed by (iii) spatiotemporal composition to learn spatiotemporal representation vector for each behavior descriptor. (iv) The model learns pre-perturbed (baseline) behavioral information as a vector form with pre-defined size. This vector is also fed into the multilayer perceptron (MLP) to learn behavior-specific perturbation responses. (v) scTRAIT transforms each perturbation into embedding vectors, which are assigned as delta vectors, and combines each with the behavioral representation vector to account for the behavioral shift due to the perturbations. (vi) All behavioral descriptors are combined and fed into the inter-behavior MLP layer to learn the association between each behavior feature. (vii) The final layer learns to predict the age or frailty status. **B**. Average accuracy score for 3 age groups classification and comparison with other models: Vanilla morphodynamic (VM) model (0.56); Vanilla trajectory (VT) model (0.76); and scTRAIT (0.84). **C**. Confusion matrix for predicting monocytes from each of the three age groups. **D**. Receiver– operating characteristic (ROC) curve of age classification (scTRAIT AUC=0.96). **E**. F1 score of age classification (scTRAIT, 0.80; VT, 0.58; VM, 0.60). **F**. ROC curve of frailty classification (scTRAIT AUC=0.97). **G**. F1 score of frailty classification (scTRAIT, 0.89; VT, 0.82; VM, 0.50). **H-I**. Pearson’s correlation heatmap between Principal components (PCs) of latent vector of the learned relationships pre and post-perturbations for age (H) and frailty (I) classifications. Symbol ● denotes correlation coefficient above 0.4.

### scTRAIT accurately classifies monocytes based on age and frailty using interpretable latent vectors

The dataset used in this analysis comprised approximately 50,000 monocytes with detailed behavioral measurements across three perturbation conditions and a control. These data were split into randomly selected training and testing subsets to evaluate the predictive performance of scTRAIT in classifying monocytes into three distinct classes based on age and frailty. Among the models evaluated, scTRAIT demonstrated the highest average accuracy score of 84%, outperforming other benchmark VT and VM models, which achieved average accuracies of 76% and 56%, respectively (**Figure 5B-C, Supplementary Figure 8A**). To further refine this approach, we investigated whether separating the prediction of age and frailty would improve the performance of scTRAIT. Rather than simultaneously predicting both age and frailty, we implemented a two-step classification process. First, the model was tasked with classifying monocytes into young versus older donors (age classification) then from the older donors, classifying old (non-frail) versus frail monocytes (frailty classification) (**Supplementary Figure 8B**). For the age classification, scTRAIT again outperformed the other benchmark models with an area under the curve (AUC) of 0.96 and an F1 score of 0.8 (**Figure 5D-E, Supplementary Figure 8C**). For the frailty classification among the older donors, scTRAIT similarly outperformed other models, achieving an AUC of 0.97 and an F1 score of 0.89 (**Figure 5F-G, Supplementary Figure 8D**). Furthermore, testing scTRAIT using technical replicates of the experiment that the model had not previously seen consistently maintained high accuracies (AUC of 0.82 and 0.91 for age and frailty classification, respectively), further establishing its reliability as a robust predictive model for age- and frailty-classifications (**Supplementary Figure 8E-F**).

While it is important to note that scTRAIT demonstrated the best overall performance, another model, VT (based on movement trajectories), exhibited comparable accuracy in frailty prediction. This observation suggests that motility alone could serve as a potential deep learning biomarker of frailty at single-cell resolution. Nevertheless, the combination of multiple behavioral modalities coupled with cellular responses to inflammatory perturbations enhances the predictive power of scTRAIT, allowing for more nuanced and reliable classification of age and frailty status.

Beyond evaluating the output prediction accuracies, we sought to investigate whether latent vectors and perturbation embeddings of scTRAIT captured biologically meaningful patterns underlying the data, rather than arising from random noise. We extracted latent representations from pre- and post-perturbation states and performed principal component analysis (PCA) to evaluate alignment with conventional human-defined behavioral features spanning 98 motility, 54 colocalization, and 12 morphodynamic features (**Supplementary Tables 2–4**). In the pre-perturbation state, the first principal component (PC0) of the motility and colocalization latent vectors showed strong correspondence with the first PC derived from their respective human-defined feature sets (**Figure 5H, I**, left), indicating that scTRAIT encoded biologically intuitive behavioral information. After perturbation, this correspondence shifted to the less variant third (PC2) and second (PC1) PCs in an age and frailty prediction, respectively (**Figure 5H, I**, right). This shift suggests that while post-perturbation responses partially capture behaviors encoded by hand-engineered features, the highest variance information was less interpretable by these traditional metrics and was better captured through more complex, non-linear representations captured by the DNN.

Visualization of the latent spaces revealed that age-associated differences were evident in the pre-perturbation state and further amplified following perturbation (**Supplementary Figure 9A, B**). In contrast, frailty-associated differences were less separable at baseline but became more distinct after perturbation, particularly within motility and colocalization latent spaces and their coupling (**Supplementary Figure 9C, D**). Analysis of perturbation embeddings showed that individual perturbations occupied distinct directions in the behavioral latent space, with IL6 exhibiting a particularly pronounced effect in the colocalization modality during age classification (**Supplementary Figure 10A, B**). In frailty classification, perturbation-induced shifts in colocalization were more prominent across DNA, IL6, and LPS treatments (**Supplementary Figure 10C, D**). Collectively, these results demonstrate that scTRAIT learns interpretable latent representations that capture both intuitive behavioral features and higher-order perturbation-dependent dynamics associated with aging and frailty.

### scTRAIT enables the development and validation of a novel cellular frailty score

Beyond classifying the age and frailty status of primary monocytes, we extended the application of scTRAIT to estimate cellular age, offering a more nuanced understanding of age-related cellular behaviors. Among young donors, we observed a correlation of 0.57 between the chronological age and predicted age, with a mean absolute error (MAE) of 3.46 years (**Figure 6A**). This strong performance relative to other benchmark models (**Supplementary Figure 11A**) highlighted a relatively linear aging trajectory in young monocytes, suggesting that the biological markers scTRAIT captures closely reflect chronological age in young donors. In the case of old monocytes, scTRAIT achieved a lower MAE of 1.74 years and a higher correlation of 0.62 between chronological and predicted ages (**Figure 6B, Supplementary Figure 11B**). Interestingly, frail cells deviated from this trend, exhibiting a plateau or similar values of the predicted age regardless of chronological age. This finding was consistent with our previous observation that frailty among older adults was an age-independent phenotype (**Figure 4D**), further reinforcing the notion that frail monocytes deviate from the linear aging process.

**Figure 6.**
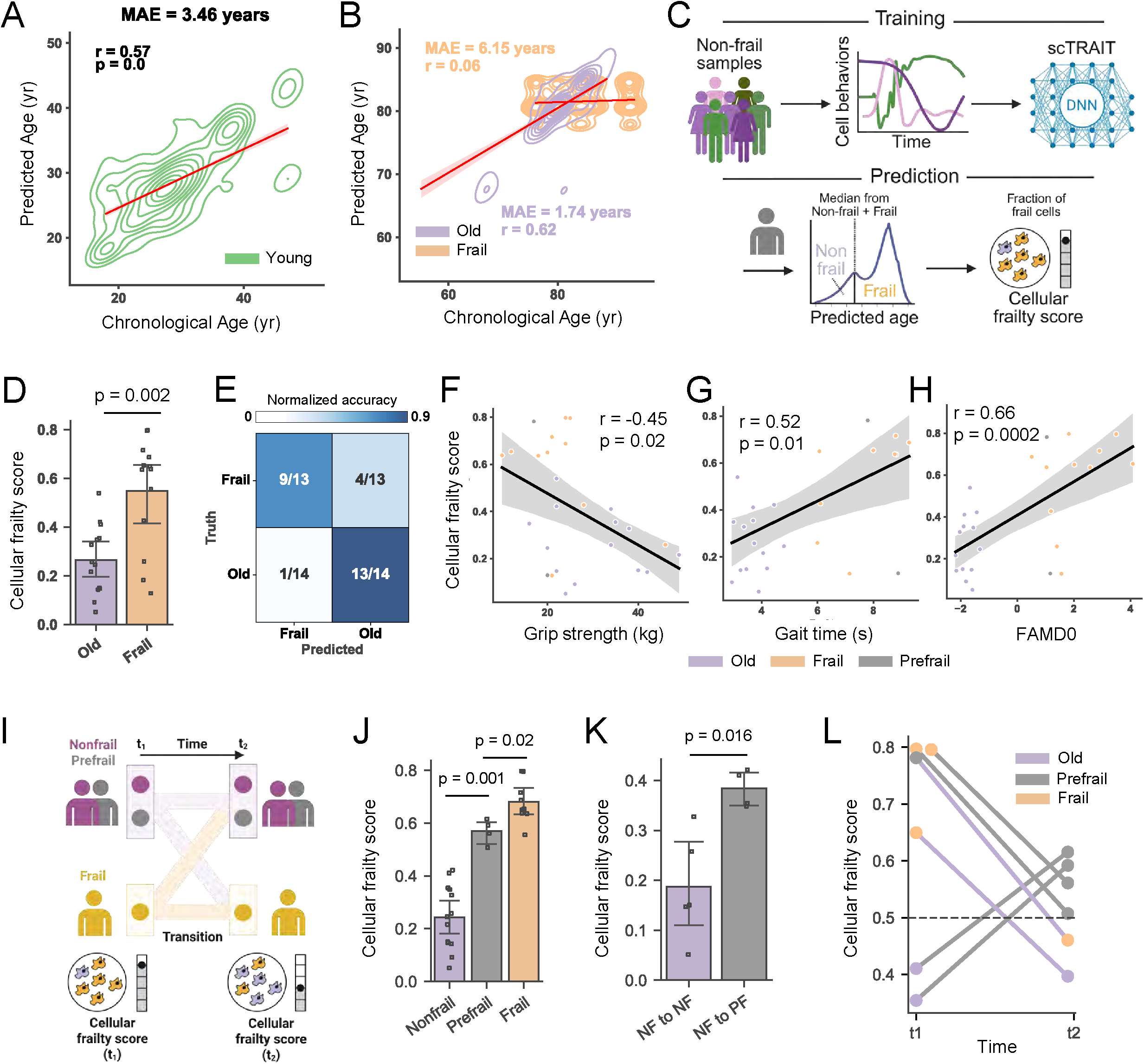
scTRAIT accurately predicts and frailty status at individual level. **A-B**. Predicted age versus chronological age across young (A) and old (B) donors. Red line denotes linear regression line with Pearson’s correlation r and means average error (MAE) shown. **C**. Schematic illustration of developing cellular frailty score (CFS) derived from scTRAIT. **D**. CFS across donors from old non-frail and older frail donors. Old donors, N= 4 donors, frail donors, N=13 donors (mean ± 95% C.I.). p-values calculated based on non-parametric Mann-Whitney U test. **E**. Normalized confusion matrix indicating the number of correct predictions based on the cellular frailty score; 9 out of 13 donors were correctly predicted as frail, and 13 out of 14 donors were correctly predicted as non-frail old. **F-H**. Regression plot between cellular and grip strength (F), gait time (G), and FAMD0 value (H). r indicates Pearson’s correlation with corresponding p-values. **I**. Schematic illustration of longitudinal analysis design. **J**. CFS across donors determined by scTRAIT for non-frail, prefrail, and frail donors (mean ± 95% C.I.). Cellular frailty score of prefrail donors were predicted to be between non-frail and frail range. Non-frail, N=13 donors, Prefrail N=4 donors, Frail N=9 donors. p-values calculated from non-parametric Mann-Whitney U test with Benjamini-Hochberg multiple test adjustment. **K**. CFS at initial time point for donors who remained non-frail and those progressing from non-frail to prefrail. NF to NF (N=5 donors) and NF to PF (N=4 donors) (mean ± 95% C.I.). p-values calculated from non-parametric Mann-Whitney U test. **L**. Prediction results based on the initial cellular frailty score for donors who naturally progressed or improved their frailty status from time t_1_ to t_2_. Period between t1 and t2 was 2 years. Colors of the dot represent the ground truth label; color of the line represents predicted label at time t_2_.

To determine the translational potential of scTRAIT, we trained an aging clock model on monocytes from old (non-frail) donors using behavior features. This was motivated by the observed decoupling between chronological age and cell behavior in frail donors. The trained model was then applied to monocytes from old and frail donors to generate donor-level distributions of predicted cellular age. Assuming that donors ultimately progress toward frailty, we quantified the fraction of monocytes that exceeded a predefined predicted-age threshold (see Methods), which was indicative of frail-like cells. This measurement was termed the cellular frailty score (CFS) (**Figure 6C**). scTRAIT successfully identified higher CFS in frail donors compared to non-frail donors that matched their clinical status (**Figure 6D**). Using a threshold of 50%, the model accurately identified most frail donors as frail and old donors as non-frail. However, four frail donors were misclassified as non-frail, and one old donor was misclassified as frail (**Figure 6E**).

To explore why scTRAIT misclassified the donors, we explored potential underlying reasons for the low CFS in the four frail donors. We investigated the post-perturbation vectors, which represented immediate frailty-associated latent vectors across different behavioral modalities. Within the motility and colocalization spaces, these donors were indistinguishable from others, indicating that these modalities did not contribute significantly to their low CFS (**Supplementary Figure 12A, B**). However, in the morphodynamic space, the misclassified donors were clearly differentiated, suggesting that the model’s predictions were largely driven by morphological plasticity (**Supplementary Figure 12C**). We questioned which perturbation responses contributed most to the low CFS in these donors. In both the motility and colocalization spaces, there was no distinctive deviation in the perturbation response of these donors (**Supplementary Figure 12D-E**, left and middle panels). However, in the morphodynamic space, particularly in response to LPS, we observed a distinct pattern that set these donors apart (**Supplementary Figure 12F**, right panel). The LPS-induced morphodynamic response appeared to contribute significantly to their lower CFS, underscoring the role of morphological changes in shaping the model’s predictions for frailty. This, in summary, scTRAIT extends beyond classification to model cellular aging trajectories and introduces CFS that accurately distinguishes the frailty status of donors based on monocyte behaviors.

### scTRAIT-based CFS encodes aspects of clinically diagnosed frailty status

To further investigate the misclassification of the four donors by our model, we sought to determine whether their CFS carried any biological or clinical significance. Specifically, we wanted to assess whether the model’s predictions were linked to measurable clinical parameters, such as grip strength and gait time (time to walk a 4-meter distance), which are key features to determine frailty using the physical frailty criteria. Remarkably, we observed significant correlations, with CFS showing correlation coefficients of -0.45 with grip strength and 0.52 with gait time, respectively (**Figure 6F, G**). These correlations of physiological measurements with the composite CFS were higher than any of the single feature-based correlations under any of the perturbations, such as average speeds (**Supplementary Figure 13A, B**). This suggests that the CFS effectively captures aspects of physical performance associated with frailty. Interestingly, the model identified some donors who were clinically categorized as frail but demonstrated relatively high grip strength and low gait times. This suggests that some clinical definitions may be insensitive to certain biological underpinnings.

In the clinic, frailty is determined by a combination of binary variables consisting of weakness, weight loss, activity levels, and exhaustion, alongside continuous variables, such as grip strength and gait time. To address this, we performed factor analysis of mixed data (FAMD) to create a composite clinical frailty feature that integrates all available measurements. When we plotted donors in a two-dimensional FAMD space, we observed a highly heterogeneous distribution among frail donors, with notable variability in their clinical characteristics (**Supplementary Figure 13C**). Interestingly, the frail donors who were predicted as non-frail by our model (due to their low CFS) tended to occupy regions of the FAMD0 axis characterized by lower values, which represented the dimension of maximum variance in the clinical data. The correlation between the CFS and the FAMD0 axis showed a correlation coefficient of 0.66 (**Figure 6H**), indicating that our CFS accurately encodes key aspects of clinical frailty features.

Taken together, the scTRAIT-derived CFS captures clinically relevant aspects of frailty that strongly correlate with traditional clinical frailty measures. However, the lack of strong correlations between clinical frailty measures and single-cell behavior features (e.g., average speeds) suggests that they are not one-to-one, and that CFS potentially captures aspects of aging and frailty that are not captured by the clinical features.

### scTRAIT-based CFS predicts and forecasts longitudinal progressions along frailty trajectories

To assess potential clinical utility and benchmark the temporal stability of scTRAIT-derived CFS captures, we conducted a longitudinal study involving a subset of 17 donors, in which frailty status was recorded at two time points: baseline (t1) and follow-up (t2). We also collected monocytes from 6 donors at t2 for single-cell behavior profiling using the scTRAIT model (**Figure 6I**). This analysis served two purposes. First, we investigated whether the CFS score obtained at t1 could prospectively forecast donor’s future frailty status at t2, that is, whether higher CFS’s at t1 were associated with an increased likelihood of transitioning towards a frail state by t2. Second, we assessed whether CFS predictions at t2 were concordant with frailty class assignments derived from the original t1 model for each donor.

Among the 17 donors included in the longitudinal analysis, 4 were clinically assessed as prefrail. Although scTRAIT did not initially include a classification for prefrail donors, we observed that the prefrail donors had intermediate scores (range: 0.5–0.62), directly positioning them between the non-frail and frail groups (**Figure 6J**). Further analysis of non-frail donors at t1 revealed a clear distinction based on their CFS. Those with relatively low scores at t1 (range: 0–0.33) maintained their non-frail status at t2, whereas those with higher CFS that were still within the non-frail range (range: 0.33–0.43) progressed to prefrail. This result suggested that higher CFS at t2 (in the non-frail range) was indicative of an increased risk of frailty progression (**Figure 6K**). Similarly, among donors initially classified as frail, those who clinically improved to a non-frail status at t2 had lower CFS values (in the frail range) compared to those who only improved to prefrail, further reinforcing the notion that the scTRAIT-derived CFS captures not only current frailty status but may be indicative of future trajectories of clinical decline (**Supplementary Figure 13D**). While these results are remarkable, further analysis using larger sample sizes is needed to systematically validate these findings.

To address our second objective, we longitudinally tracked frailty trajectories based on the behaviors of monocytes from each donor. Of the six donors evaluated, scTRAIT correctly classified five donors at t_2_ based on their CFS. Among these, two donors improved from frail to prefrail, one donor transitioned from frail to non-frail, and two donors moved from non-frail to prefrail (**Figure 6L**). Importantly, the alignment between clinical frailty status and CFS classification was further supported by the observed changes in CFS over time. Donors who progressed to a prefrail state exhibited increased CFS, whereas those who improved from frail to non-frail or prefrail showed corresponding decreases in their CFS.

Collectively, these results highlight the robustness and translational relevance of the scTRAIT-derived CFS as a sensitive marker of both current physiological frailty and future risk trajectories. Our approach captures cross-sectional frailty status and additionally reflects longitudinal shifts, reinforcing its potential as a sensitive and dynamic indicator of aging and frailty trajectories in humans.

## DISCUSSION

Our study presents significant advancements in understanding the relationship between single-cell behaviors and the processes of aging and frailty. We identified associations of age and frailty with distinct single-cell motility patterns, responses to inflammatory perturbations, and divergent effects of local cell density, highlighting the multifactorial regulation of monocyte behaviors. Monocytes are highly adaptable innate immune cells that exhibit both intrinsically encoded and extrinsically induced states^45^. With aging, monocytes exhibit well-described functional changes, including mitochondrial dysfunction, reduced phagocytosis, impaired antigen presentation, altered chemotaxis, increased basal inflammatory signaling, and changes in tissue-resident phenotypes, which can collectively contribute to inflammaging. Inflammaging represents a milieu for multisystem dysfunctions and arises from diverse molecular mechanisms, including genetic susceptibility, DNA methylation remodeling, metabolic reprogramming, and inflammatory proteomic changes, which together contribute to frailty phenotypes in humans^29,46^. These age-associated molecular shifts in monocytes are, in part, expressed through heterogeneous behavioral phenotypes, reflecting distinct inflammaging pathways that contribute to age- and frailty-related systemic and tissue dysfunctions.

The development of the scTRAIT model represents a significant methodological advancement that is different, yet complementary, to recent advances in aging and frailty research. By combining spatiotemporal descriptions of cell behaviors and responses to perturbations, scTRAIT effectively predicts age and frailty, making it a valuable tool for future research and potential clinical applications. scTRAIT aligns with current efforts to quantify biological age and identify a range of biomarkers known as aging clocks. At the molecular level, omics-based measures that focus on age-related changes in epigenomics, transcriptomics, proteomics, and metabolomics provide important insights into the mechanisms of aging^47^. For example, various aging clocks have been developed based on DNA methylation at CpG sites, which could reflect stress accumulation and replication errors^48^. Cell behaviors represent the combined effects of age- and frailty-related dysregulation of core biological functions^49–51^. Thus, scTRAIT functions, in part, as a cell behavior-based aging clock that quantifies cellular function and adaptive capacity in real time, providing relevant features for defining modern clinical definitions of aging and frailty.

While scTRAIT demonstrates strong predictive capability in identifying frailty based on monocyte behaviors, the biological meaning of the CFS will require further study, ideally using both cross-sectional and longitudinal donor samples. Furthermore, directly correlating the CFS, which captures cell behavioral nuances, with underlying molecular profiles could provide deep mechanistic understanding and the identification of novel therapeutic targets. Future research should focus on this to identify specific molecular markers, such as inflammatory cytokines, epigenetic modifications, metabolic signatures, or differentiation patterns, which should be simultaneously assessed with single-cell behavior patterns. In fact, as circulating monocytes can serve as precursors to tissue-resident macrophages, frail monocytes may give rise to dysfunctional macrophage populations that contribute to impaired tissue homeostasis, chronic low-grade inflammation, and diminished regenerative capacity, which are hallmarks of frailty^52,53^. Longitudinal and functional assays will be required to clearly understand these differentiation pathways, which could also inform therapeutic strategies aimed at selectively targeting or reprogramming frail monocyte behavioral states before entering peripheral tissues. Linking the cell behavior to these molecular underpinnings could lead to a shift in how frailty is clinically defined, moving from a primarily physiological and functional assessment toward a more nuanced, biologically informed framework. This research could provide new strategies to determine whether eliminating frail monocytes influences the progression of frailty at both physiological and clinical levels.

Quantifying frailty at the single-cell level offers a new framework for refining current paradigms to classify frailty into subtypes with distinct clinical trajectories and biological drivers. Clinically defined frailty subtypes characterized by mobility impairment, exhaustion, and weight loss were associated with different long-term outcomes and progression to multimorbidity, thus underscoring that frailty does not necessarily arise from a single uniform mechanism ^54^. Molecularly defined frailty subtypes, including metabolomic and inflammatory profiling, have revealed subtype-specific signatures linked to chronic inflammation, metabolic dysregulation, and systemic physiological decline, leading to differential responses to lifestyle or therapeutic interventions^55,56^. Furthermore, our single-cell approach to characterizing frailty could reveal new subtypes of frailty, or subpopulations of individuals who may be susceptible to frailty progression, or potentially those who could maximally benefit from certain interventions.

Lastly, our behavior-based frailty framework offers a complementary approach to addressing this heterogeneity by capturing meso-scale functional characteristics of underlying mechanisms at the single-cell level, all while ensuring experimental simplicity and scalability. Moreover, the non-invasive nature of image-based single-cell behavior analysis makes it an appealing method for clinical applications. It can facilitate early detection, track and forecast longitudinal age-related decline and frailty progression, and identify targetable strategies and interventions to reprogram dysfunctional cell behaviors into healthier states, ultimately improving health outcomes in aging populations. Altogether, our findings reveal that single-cell behaviors of monocytes robustly encode information about aging and frailty, establishing monocytes as significant biological sensors of aging and frailty in humans.

## MATERIALS AND METHODS

### Recruitment of primary donor samples

All donors were recruited through the Johns Hopkins Older Americans Independence Center (OAIC) registry by well-trained clinical research staff in accordance with approved IRB protocols. Briefly, donors fitting the criteria of age and frailty status were recruited to Johns Hopkins Hospital at Bayview, and whole blood was drawn by a trained phlebotomist or clinician. From the blood, we isolated peripheral blood mononuclear cells (PBMCs) using Ficoll-based density centrifugation, and monocytes were further isolated from the PBMCs (see section below). Ground truth labels of frailty among the older donor samples were determined based on the Physical Frailty score^57^. In our study, non-frail/robust older adults had a score of 0, prefrail donors had a score of 1 or 2, and frail donors had scores of 3-5.

### Monocyte isolation and cell culture

Human PBMCs were isolated from the peripheral blood of young, old (non-frail), prefrail, and frail donors. Cells were plated at a culture density of 1 × 10^6^ cells/mL in RPMI base medium that contained 1× Glutamine (Gibco) and was supplemented with 10% heat-inactivated fetal bovine serum (FBS), 5% Penicillin/Streptomycin, and 5% HEPES (1M). Primary CD14^+^/CD16^-^ monocytes were further purified from other PBMCs using an EasySep human monocyte immunomagnetic isolation kit (STEMCELL Technologies) and EasySep Magnet (STEMCELL Technologies) according to the manufacturer’s protocol. Monocytes were then centrifuged at 350 g for 5 min and resuspended in complete RPMI (described above) at a concentration of 2.5 × 10^5^ cells/mL. In experiments involving cell tracking, monocytes were labeled with 0.01mM CellTracker Green fluorescent dye (ThermoFisher Scientific) for 20 min and subsequently washed to enable the downstream cell tracking.

### Cell seeding and exposure to *ex vivo* perturbation/stressors

Rat tail collagen type-I was neutralized and diluted to 1 mg/mL with the addition of 0.2 M NaOH, 1× Hanks Balanced Salt Solution (HBSS), and complete RPMI^58–60^. All reagents were mixed thoroughly on ice for 1-2 min to ensure uniformity and a pH of 7-7.5. Subsequent dilution was made to a final working concentration of 0.5 mg/mL by mixing equal parts of 1 mg/mL collagen and the monocyte cell suspension media to obtain a final monocyte concentration of 2.5 × 10^5^ cells/mL of gel. We transferred 100 µL of the monocyte-containing gel mixture to each well of a 96-well glass-bottom plate (Cellvis), ensuring uniform coverage without cell aggregation at the walls. Once the monocyte-laden gel precursors were loaded into each well, the 96-well plate was maintained at 4°C for 10-15 min to inhibit gelation and allow monocytes to sediment to the bottom of the wells. Following this 5-10-minute settling step, the plate was transferred to a 37°C incubator for 1 hour to allow gelation of the collagen, thereby immobilizing monocytes just beneath the soft collagen gel. After complete collagen polymerization, all monocyte-containing gels were gently hydrated by adding 100 µL of complete RPMI on the top of the gels. In perturbation experiments, wells were alternatively hydrated using 100 µL of pre-warmed complete RPMI supplemented with the proinflammatory agents, including either 100 ng/mL recombinant human interleukin-6 protein (IL-6; R&D Systems), 10 µg/mL lipopolysaccharides from Escherichia coli (LPS; Sigma Aldrich), or 1µg/mL poly(dA-dT) synthetic double-stranded free DNA (Invitrogen) for 1 day prior to imaging.

### Live-cell imaging

Time-lapse brightfield and fluorescence images of monocytes were acquired every 2 min for up to 3 hours using a Leica Stellaris 5 confocal microscope at 20X magnification (0.568 um/pixel),1024×1024 pixel^2^ scanning resolution, and an acquisition speed of 600 Hz. All cells were imaged while encased in a 5% CO_2_-regulated chamber at 37°C. Focus planes were determined from the bottommost layer of cells just above glass substrates.

#### Image and trajectory processing

For processing the live images, we developed scDynamics built on the in-house CaMI framework^61^ which automatically detects, and measures single-cell behaviors followed by unsupervised identification of behavior clusters. This automatic method enabled us to avoid human bias and increase sample sizes to gain statistical power. Our optimized workflow sequentially performs 1) single-cell segmentation, 2) tracking and trajectory registration, 3) feature extraction, 4) dimensionality reduction and unsupervised clustering identification. Each of these steps is described below.

#### 1) Single-cell segmentation and tracking

Masks of the single-cell monocytes were identified using a deep neural network-based generalist algorithm for segmentation: Cellpose 1.0 using the cyto2 model^62^. Thresholding and segmentation diameters were evaluated and optimized for each experiment to remove any detected cell debris. The centroid locations of the generated single-cell masks were used for linking the cellular entities in subsequent frames to perform single-cell tracking over successive time intervals. To achieve this, we used the Python package Trackpy 0.5.0, which is based on the Crocker-Grier algorithm^63^, and an adaptive search range was used to enhance the quality of tracking followed by visual inspection of the trajectories. We discarded any trajectory that skipped at least one frame to improve tracking accuracy. At every instance we recorded centroid locations, multiple morphology features, and spatial colocalization for each trajectory using our in-house feature quantification pipeline.

#### 2) Trajectory registration

To avoid effects related to variations in the length and direction of trajectories, we temporally registered all trajectories to a length of 1h. Trajectories that were tracked for less than 1h were discarded; those tracked for more than 2h were segmented into multiple trajectories to retain highly qualified tracks and to maximize the sample size. We wanted to ensure independence of the directions in which cells move, which is an external factor varied by setting the region of interest; therefore, we defined the primary migration axis based on the maximum length of displacement measured throughout the observation period, with a nonprimary migration axis that is orthogonal^64,65^. To obtain these axes, we applied singular vector decomposition (SVD) to the displacement matrix of cells *A*.

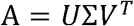

Here, U is a matrix of eigenvectors of AA^*T*^, *V*^*T*^ is a matrix of eigenvectors of A^*T*^*A*, and ∑ is a diagonal matrix of singular values. Matrix multiplication is applied on the matrix of coordinates *R* with *V* to get *R*_*reg*_ = *RV*. The first and second column of *R*_*reg*_ describe the movement paths along the primary and nonprimary axes, respectively. Also, all trajectories are translated to set the initial point as the origin.

#### 3) Behavior feature extraction

We chose 98 motility and 12 morphodynamic features to describe single-cell behaviors of monocytes. Motility features include displacement-based features and turning angle-based features that capture magnitude, distribution descriptors, signal descriptors, temporal correlation, entropy, and decomposed features^66,67^. A detailed list of motility features is described in Supplementary Table 2. For parametrizing morphodynamics, we constructed a two-dimensional morphology PCA space from 15 morphology features including cell area, perimeter, convex area, solidity, eccentricity, equivalent diameter, extent, major axis length, minor axis length, aspect ratio, elongation, compactness, roundness, circularity, and rectangularity. For each movement trajectory, a corresponding ‘morpho-trajectory’ that traverses the two-dimensional morphology space can be recorded to quantify features that are associated with displacement, turning angle, distribution descriptors, and autocorrelation. A list of morphodynamic features is described in Supplementary Table 4.

#### 4) Dimensionality reduction and unsupervised clustering

We conducted PCA on multidimensional feature analyses to remove covariances between the parameters and used PCs of 95% cumulative variance to generate two-dimensional UMAP projections^68^. We used K-means ++ unsupervised clustering algorithm to define nine MCs from PCs. To validate the clustering, we projected the trajectories from each MC to qualitatively evaluate similarities among the trajectories. To quantify motility state heterogeneity, we calculated the Shannon entropy *S*.

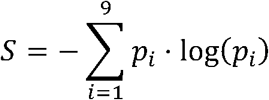

Here *p*_*i*_ is the fraction of cells within each motility cluster *i*.

#### Spatial localization analysis

For local neighbor detection, we used a search range of 100 μm and quantified the number of cells within the boundary for each frame to obtain time series of neighbor counts. “Isolated” and “close” conditions were determined based on the average number of neighbors to be above 9 or below 0.6 throughout tracked duration, respectively. For single cell-to-cell proximity analysis, we generated the matrix of centroid distances for all pairs of cells in the region of interest (ROI) for each frame to obtain time series of single-cell shortest distance, and time series of average distance from the neighboring group of cells. We also extracted 54 colocalization features from the time series of neighbor size, shortest distance, and average group distance to characterize dynamic local cell density including average, coefficient of variance (CV), slope of neighbor size, and directional bias towards the neighboring cell. A detailed list of colocalization features is provided in Supplementary Table 3.

#### Quantifying age axis coefficient (AAC)

We adopted a simple framework that captures the linear relationship between cellular motility behaviors and chronological age across a population^38^. Given a set of motility features that creates a multi-dimensional motility space, we computed centroids of cell populations for each donor. We then established a linear age axis by using the youngest donor in the cohort as the initial point, and oldest as the terminal point for each age group. The AAC is calculated by projecting each donor’s cellular behavior onto this linear age axis by following formula:

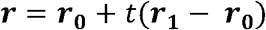

Here ***r***_**0**_ is a centroid vector from the motility space of the youngest donor, ***r***_**1**_ is a centroid vector of the oldest donor, ***r*** is a constructed linear age axis, and *t* is AAC. AAC quantifies the alignment between a donor’s motility features and the linear trajectory of aging to reflect how closely a donor’s cellular behavior corresponds to the expected pattern of cellular aging. Higher coefficients indicate closer alignment with the behaviors of older donors, and lower coefficients suggest younger cellular behaviors.

#### scTRAIT

Given a dataset of *N* cells 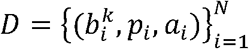, where 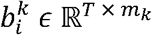 is a *T* length sequence of *m*_*k*_ dimensional behavior of cell *i* with behavior modality *k*. For simplicity, we fixed the number of modalities to three, and corresponding dimensions, where *m*_*k*=1_ = 2, *m*_*k*=2_ = 4, *m*_*k*=2_ = 8. The dataset also contains *p*_*i*_ *ϵ* ℤ describing the indexes of the perturbation applied on cell *i*, and *a*_*i*_ *ϵ ℝ* corresponds to age and frailty status, which is the target outcome of the scTRAIT to learn mapping function *f*_*scTR*A*IT*_ given behavior 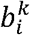, and perturbation *p*_*i*_. Below we describe the four core parts of scTRAIT.

#### 1) Behavior encoder

The first step of the model is to learn spatiotemporal encoder 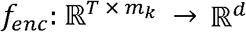 to produce latent representation 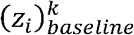 that encodes baseline spatiotemporal characteristics of the behavior.

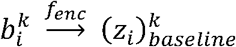

We sought to design the encoder in a way that separately learns spatial characteristics in *m*_*k*_ dimensions and temporal characteristics in *T* dimensions by learning spatial encoder 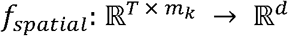 and temporal encoder 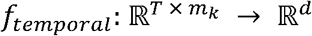 to learn latent spatial vectors 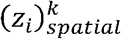 and 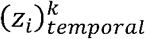 respectively^69,70^. The spatial encoder consists of series of 2-dimensional residual convolution blocks followed by dense layer and leaky ReLU activation, and the temporal encoder applies the temporal convolutional network with respective dilation rate of 1, 2, 4.

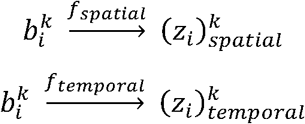

Next, in the spatiotemporal composition these two latent vectors are concatenated, and a single dense layer with weights *W* and bias *b* followed by leaky ReLU activation *σ* is applied to extract the baseline latent vector 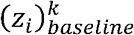.

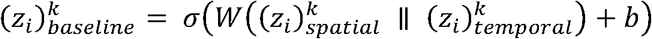

where ∥ represents vector concatenation.

#### 2) Behavior-specific perturbational deviation

scTRAIT incorporates the perturbation responses^71^ specific to behavior and behavior modality and learns the mapping 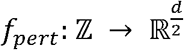 to get post perturbation behaviors 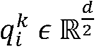 from perturbation indexes *p*_*i*_.

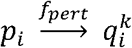

Perturbation indexes *p*_*i*_ *ϵ* ℤ are transformed from scalar form to perturbation latent embeddings *e*_*i*_ *ϵ ℝ*^*d*^ via embedding layer. To learn behavior-specific perturbational deviation 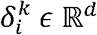, we combined both the perturbation embedding and baseline latent behavior to apply a multi-layer perceptron (MLP).

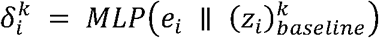

Our MLP consists of three dense layers followed by leaky ReLU activation and dropout of 0.1 in between. The model calculates the post perturbed behavior 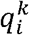 by performing element-wise addition followed by MLP.

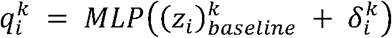

#### 3) Age decoder

The final step of the model is to decode the behavior information to age and frailty status. The model learns the association across the behavior modalities to map it to a combined behavior vector 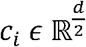.

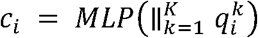

where *K* is the size of the behavior modalities. Lastly, the model results in the predicted age and frailty status 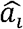 by applying MLP and additional dense layers on *c*_*i*_.

#### 4) Optimization and loss function

scTRAIT optimizes trainable parameters using Adam optimizer with a learning rate of 0.001. The loss function varies based on the objective that the model pursues. We used binary cross entropy *L*_*BCE*_, sparse categorical entropy *L*_*SCE*_, and mean absolute error *MAE* for binary classification, multi-class classification with the number of classes C, and age regression, respectively.

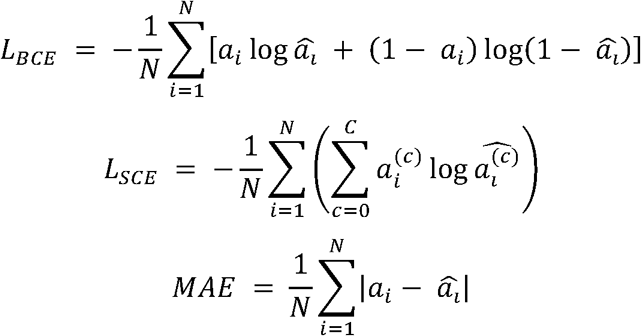

#### Benchmarking models

Two benchmark models were built to evaluate how incorporating multiple behavior modalities with the perturbation responses enhances the performance of the model. The baseline trajectory model (VT) and baseline morphodynamics model (VM), each inputting either motility or morphodynamics respectively, were designed with architecture similar to that of scTRAIT, which comprises a behavior encoder followed by an age decoder, but is absent of a behavior-specific delta layer and an inter-behavior association layer to remove any model structure effect.

#### Model evaluation

The performance of the model was quantified based on the accuracy, precision, recall, F1 score, and the area under the curve (AUC) of the receiver operating characteristic (ROC) curve.

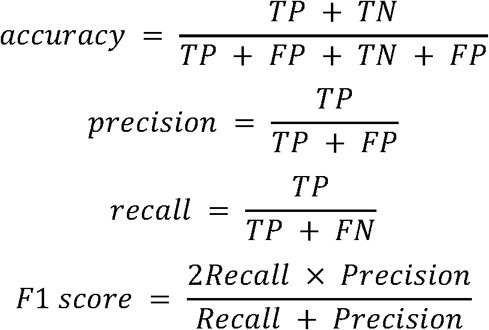

#### Cellular frailty score (CFS)

We trained the model using the cells from the age spectrum of non-frail donors and predicted with each donor a leave-one-out validation approach to ensure that each donor predicted is new data that the model has not seen previously. The outcome of each donor prediction is a distribution of predicted age, with a non-frail and frail old aggregate median value of 80. We defined the CFS as a fraction of cells that are above the median of the predicted age distribution, which represents the fraction of ‘frail’ phenotype cells.

#### Factor analysis on mixed data (FAMD) on clinical frailty determinants

To analyze the clinical and behavioral data, we utilized FAMD, a statistical method designed to integrate both continuous and categorical variables simultaneously. This approach allowed us to integrate diverse clinical measurements, including binary variables, which include weakness, weight loss, activity levels, and exhaustion alongside numerical variables including grip strength and gait time. FAMD identifies key dimensions that capture the maximum variance in the combined dataset, providing a low-dimensional representation of the clinical frailty landscape. By projecting donors into this basis, we assessed the relationships between their clinical characteristics and frailty status.

#### Statistics and reproducibility

To establish statistical comparisons between multiple groups in Figure 1, we performed the Kruskal-Wallis test followed by Dunn’s post-hoc test. Multiple statistical comparisons between the control and perturbation in Figure 2 were based on a One-way ANOVA test followed by Dunnett’s post-hoc test. For all statistical comparisons between two groups, we used the nonparametric Mann-Whitney U test. Correlations of linear regression fits were evaluated using r and p-values, respectively. Data was scaled, where applicable, to normalize distributions for gaussian-based models. No data was excluded from this work. The experiments were not randomized, and the authors were not blinded to the age- or frailty-specific demographics.

#### Software

All single cell segmentations were performed using optimized workflows in CellPose. Behavior feature extraction was performed using a custom algorithm combining CellPose outputs with TrackPy algorithm. All analyses were performed in Python with the following software specifications: python 3.9.13, tensorflow 2.10.1, scipy 1.13.1, scikit-learn 1.1.13, scikit-image 0.19.3, scikit-posthocs 0.9.0, statsmodels 0.13.5, pandas 1.5.2, matplotlib 3.6.2, seaborn 0.11.2, trackpy 0.5.0, umap-learn 0.5.3, numpy 1.23.5, cmcrameri 1.9, EntropyHub 0.2, and pyarrow 12.0.1.

## Code availability

The source code used to generate the data and findings presented in this work is deposited on the Phillip Lab GitHub page https://github.com/Phillip-Lab-JHU/scDynamics.

## Data availability

All data supporting the findings of this study are available within the paper, its Supplementary Information, and the Phillip lab GitHub page https://github.com/Phillip-Lab-JHU/scDynamics.

## Acknowledgements

We acknowledge the financial support for this work from The Johns Hopkins University Older Americans Independence Center (OAIC) pilot award through the National Institute on Aging under award number P30AG021334 (JMP); R35GM157099 from the National Institute for General Medical Sciences (JMP); Catalyst Award from the National Academy of Medicine through the Global Healthy Longevity Challenge (JMP); and the 2024 Salisbury Family and Center for Innovative Medicine Human Aging Project Scholar Award (JMP).

The following figures were created using Biorender: Figures 1A, 3A, 5A, 6C and 6I

## Author contributions

CM, PA, JW, EJP, and JMP conceived and designed study; CE, YWD, NM, and LT performed experiments; LN, AK, CS, JL, PA, and JW coordinated and collected primary donor samples; CM and JMP conceived and developed prediction models and performed formal analysis; CM, PA, JW, EJP, and JMP interpreted results; CM, EJP, PA, and JMP wrote the manuscript; All authors edited and contributed to the final manuscript

## FIGURE CAPTIONS

**Supplementary Figure 1.**
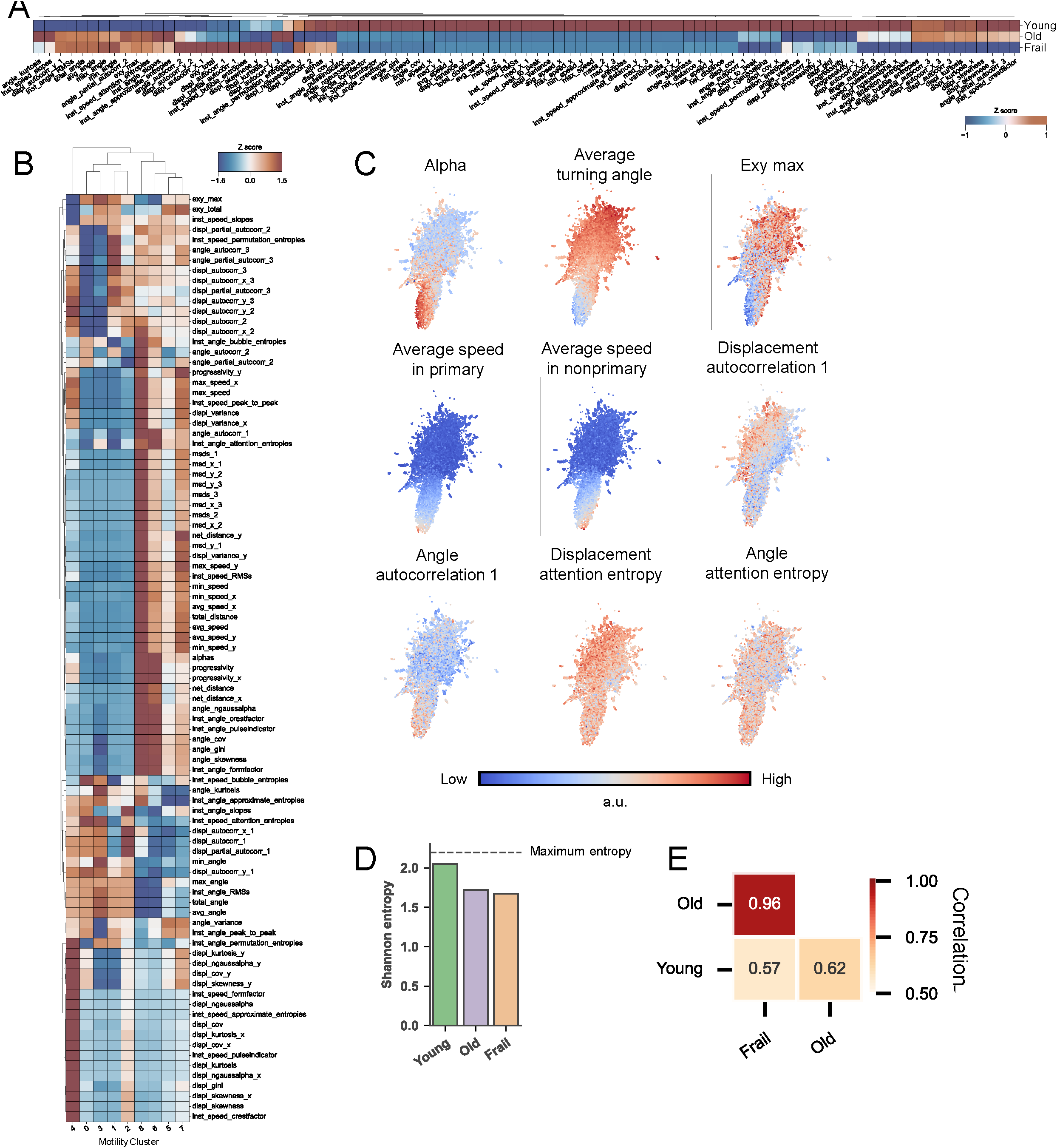
**A**. Z scores across motility parameters for each age group. Hierarchical clustering is based on the Euclidean distance using Ward method groups similar Z scores of features. **B**. Z scores across motility parameters for each motility cluster. Hierarchical clustering is based on the Euclidean distance using Ward method groups similar Z scores of features. **C**. Projection of various motility features onto UMAP space. **D**. Shannon entropy to quantify heterogeneity of motility states. The dotted line indicates the maximum entropy that can be calculated from nine motility cluster distributions. E. Cross correlation based on the fraction of motility clusters.

**Supplementary Figure 2.**
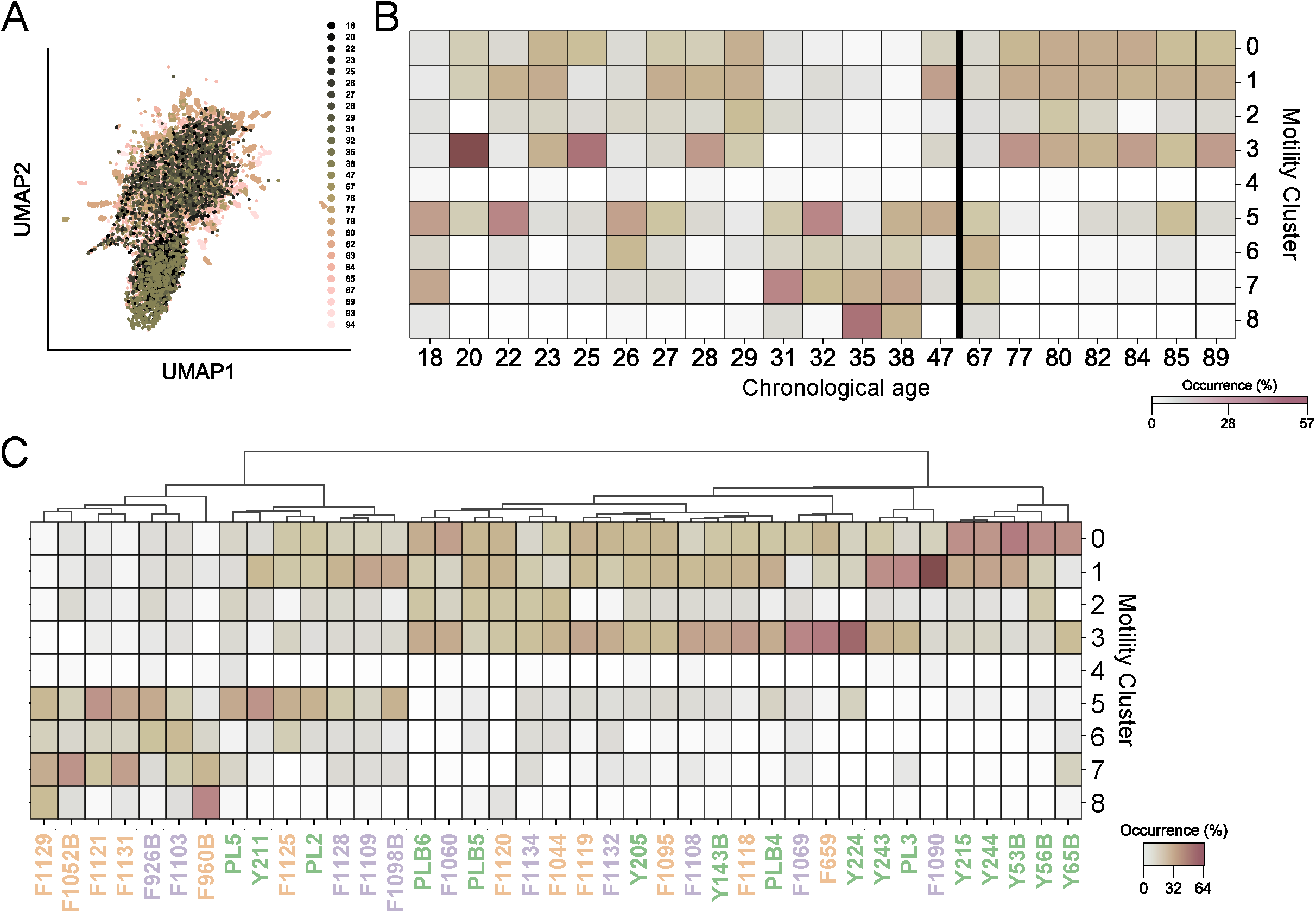
**A**. Spectrum of age projected onto the UMAP space describing the donor-to-donor variation within the young, old and frail groups for control condition. **B**. Motility cluster distribution for each age under control, suggesting non-linear and heterogeneous cluster enrichment patterns especially within the young. A bold line distinguishes the young and old group. **C**. MC distribution across all donors in control condition highlights the donor-to-donor variation in the motility patterns within each age group. Green, purple, orange colors represent young, old, frail groups respectively.

**Supplementary Figure 3.**
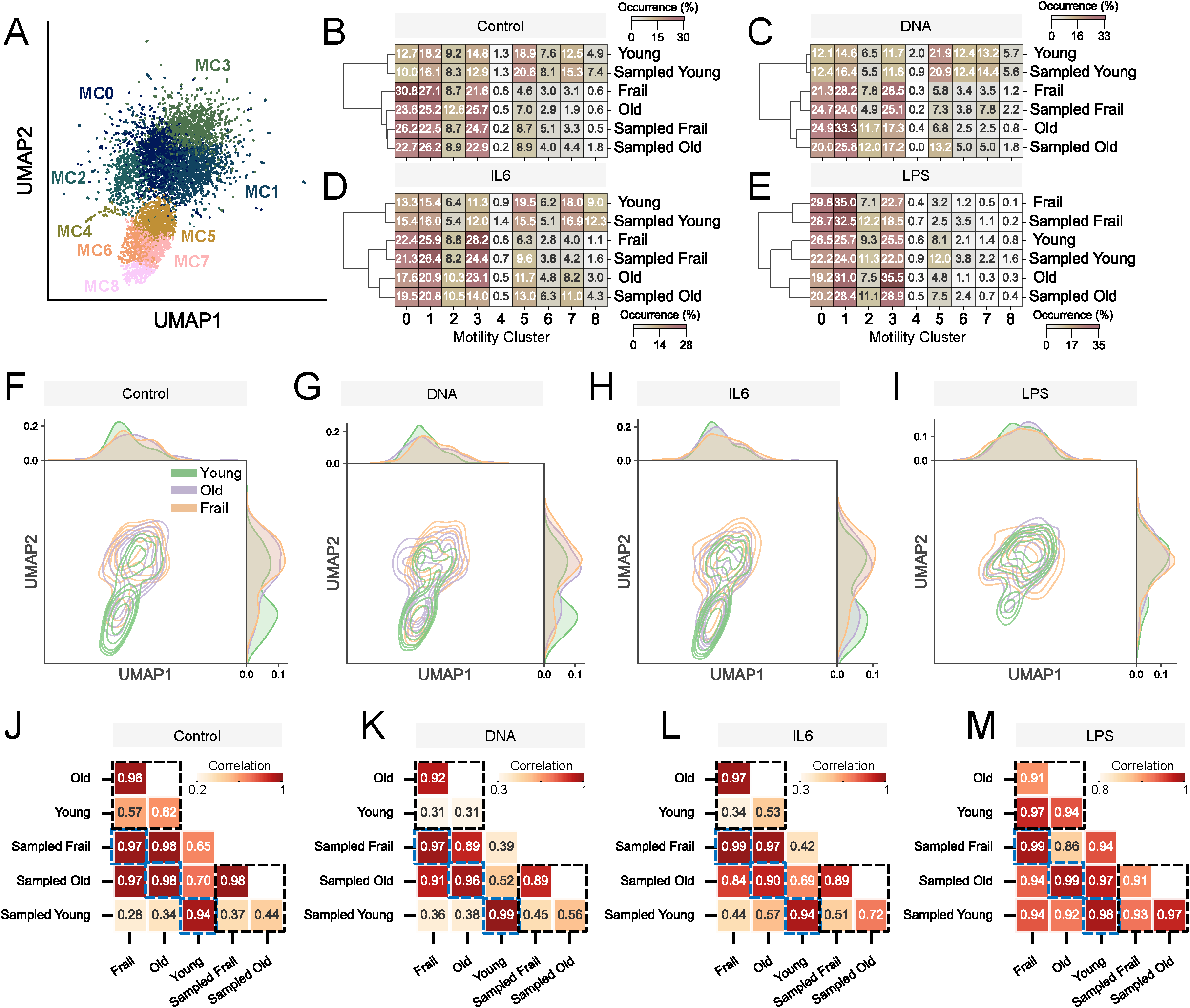
**A**. 2D UMAP representation of nine motility clusters in the sampled space (sampling n = 50 each donor). **B-E**. Heatmap showing the fractional occurrence of nine MCs across the age groups and sampled age groups exposed to control (B), DNA (C), IL6 (D), and LPS (E). Hierarchical clustering is based on the Euclidean distance using the Ward method. The sum of rows equal 1. **F-I**. 2D UMAP KDE representation of sampled space across age groups exposed to control (F), DNA (G), IL6 (H), and LPS (I) conditions. 1D distributions of single cells for each at the top and right margins. **J-M**. Cross correlation based on the fraction of motility clusters exposed to control (J), DNA (K), IL6 (L), and LPS (M). Top left and bottom right dotted black box represent comparison within original and sampled space, respectively. Blue dotted box represents comparison between original and sampled space for each age group.

**Supplementary Figure 4.**
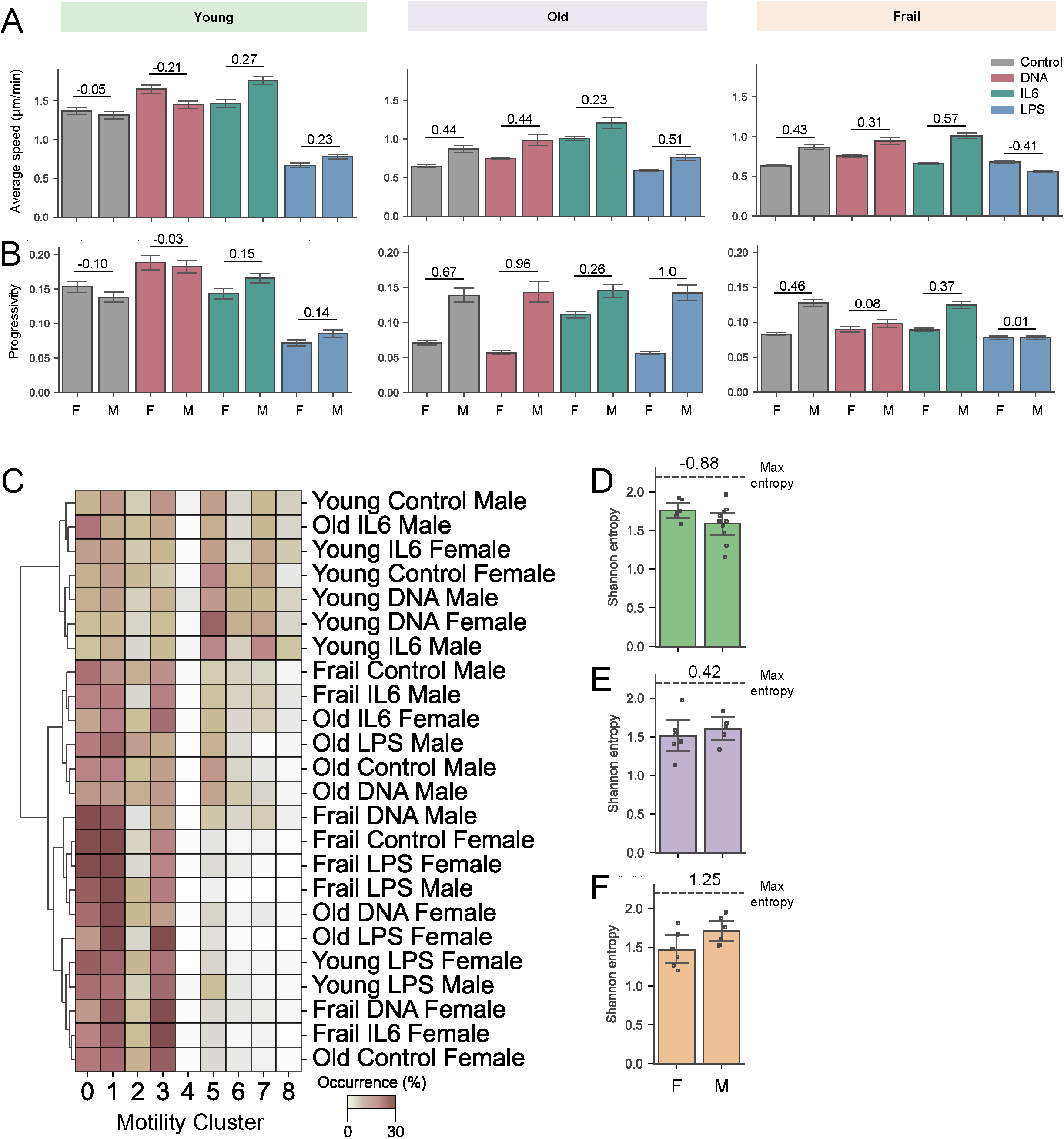
**A-B**. Bar plot comparing female (F) and male (M) with respect to control, DNA, IL6, and LPS based on average speed (A), and progressivity (B). Annotated values represent Cohen’s d. **C**. MC distribution of cells that are categorized combining age groups, perturbation and gender. Hierarchical clustering is based on the Euclidean distance using Ward method. **D-F**. Shannon entropy in control group comparing female (F) and male (M) within young (D), old (E) and frail (F) group (mean ± 95% C.I.). Annotated values represent Cohen’s d (young female N=6, young male N=10, old female N=6, old male N=5, frail female N=6, frail male N=6 donors).

**Supplementary Figure 5.**
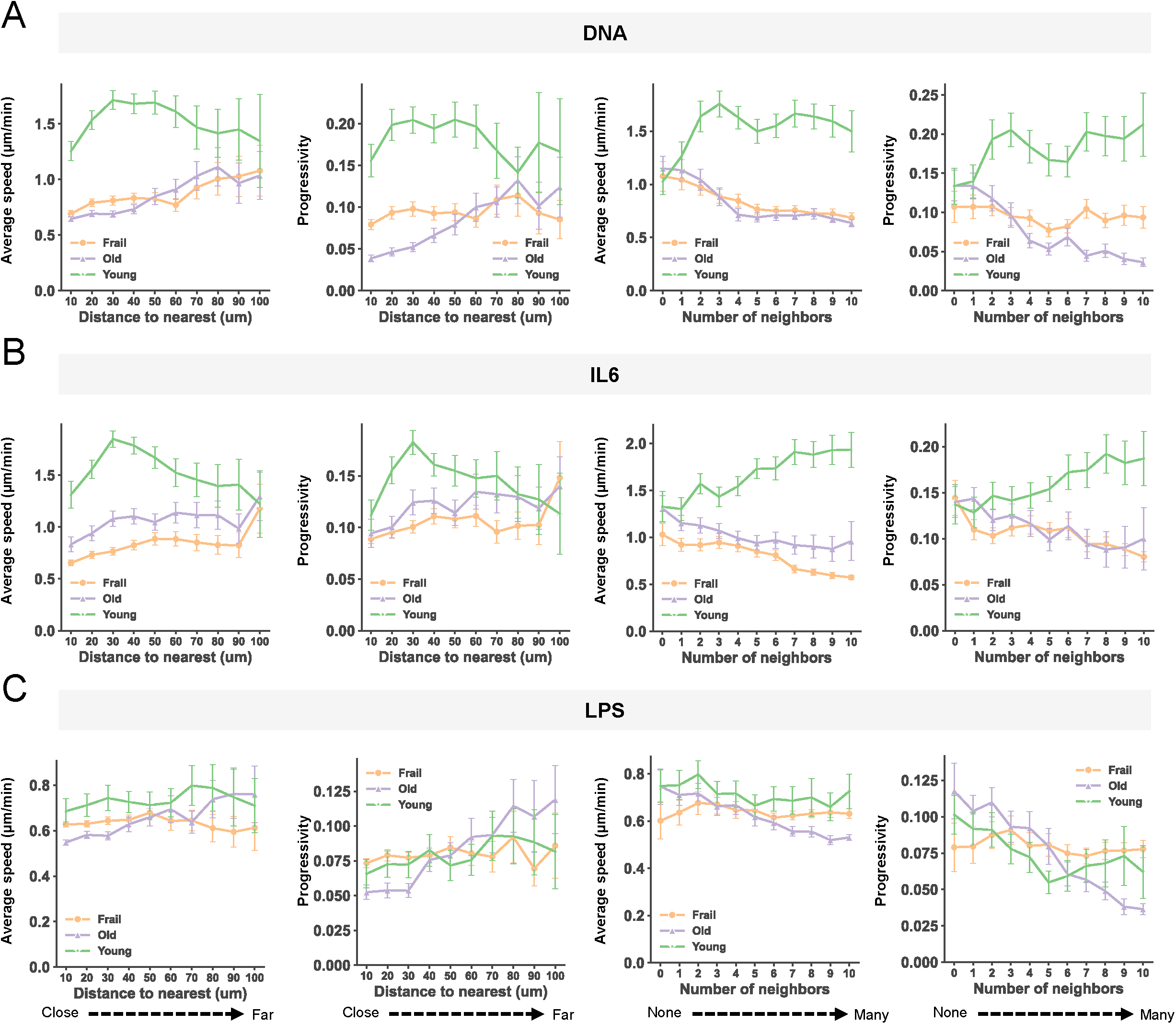
**A-C**. Average speed and progressivity over the distance to the average nearest cell with 10 µm bin (mean ± 95% C.I.) and over the average number of neighbors within the 100 µm range with 1 cell bin (mean ± 95% C.I.) under DNA (A), IL6 (B), LPS (C) conditions.

**Supplementary Figure 6.**
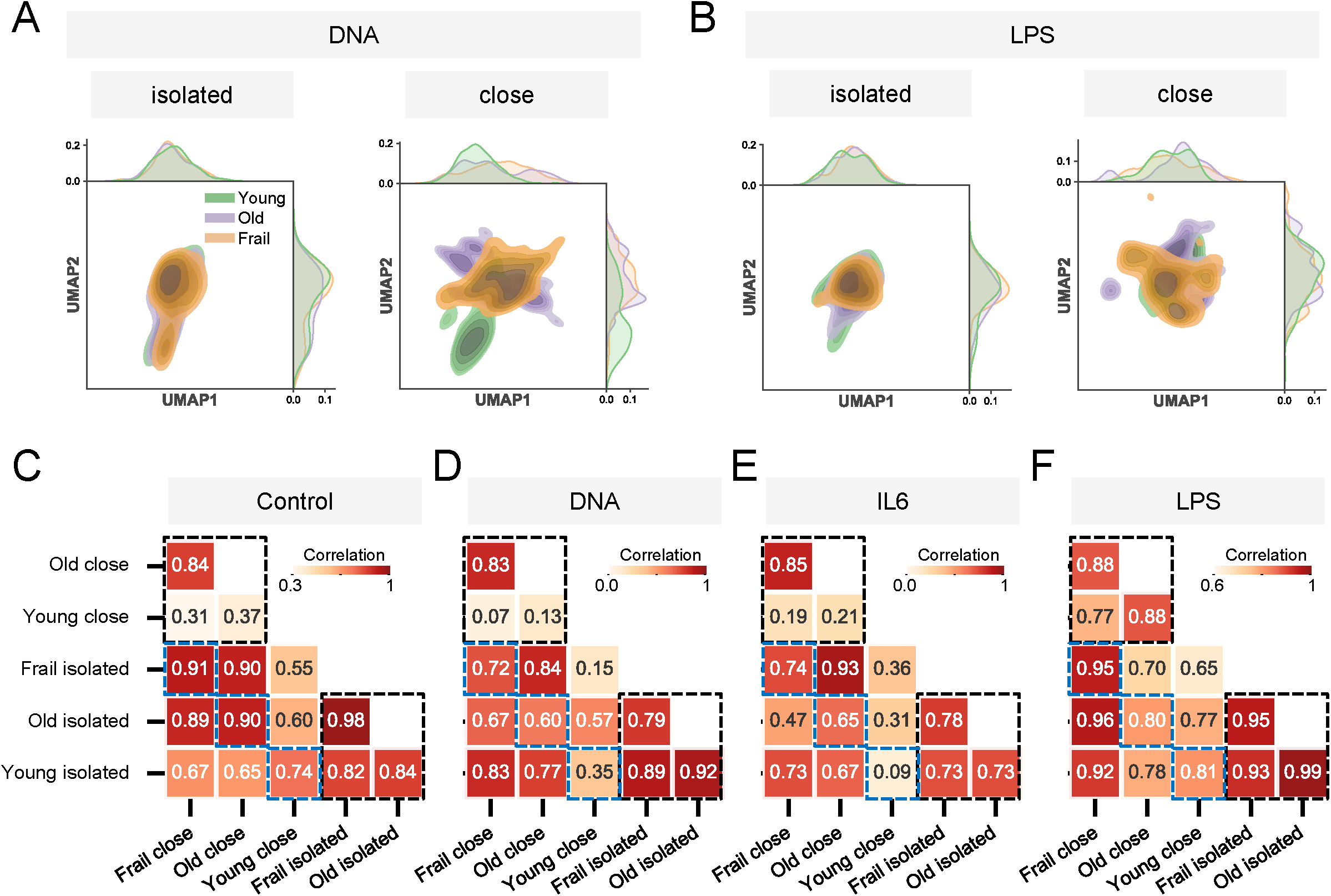
**A-B**. UMAP projection of motility distribution comparing isolated and close conditions within DNA (A) and LPS (B) treated experiments. **C-F**. Cross correlation based on the fraction of motility clusters for 3 age groups with each isolated and close condition within Control (C), DNA (D), IL6 (E), and LPS (F) treated samples. Top left and bottom right dotted black box represent comparison within close and isolated monocytes, respectively. Blue dotted box represents comparison between close and isolated cells for each age group.

**Supplementary Figure 7.**
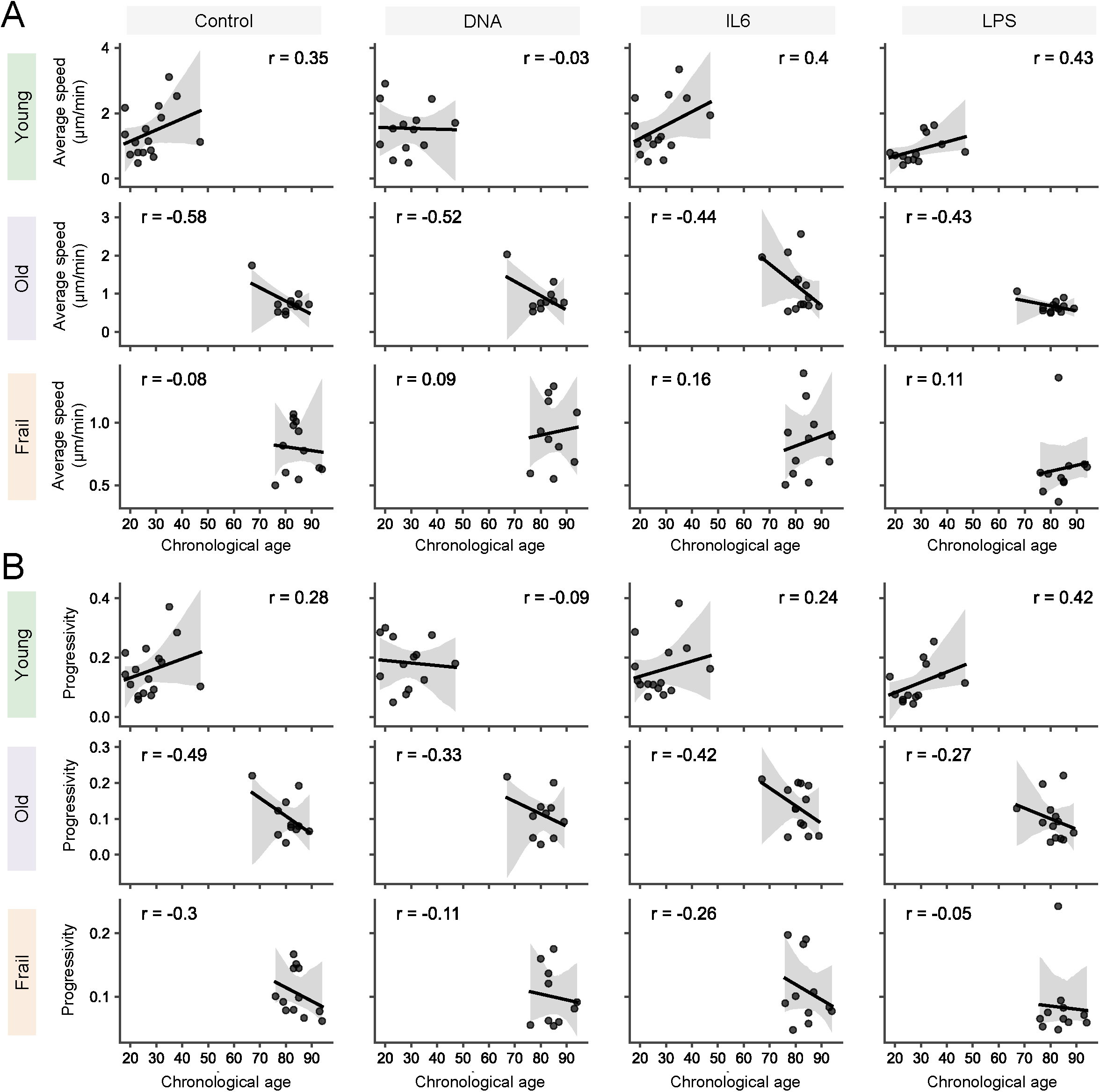
**A-B**. Regression plot of average speed (A) and progressivity (B) over chronological age across 3 age groups. Annotated r refers to Pearson’s correlation coefficient.

**Supplementary Figure 8.**
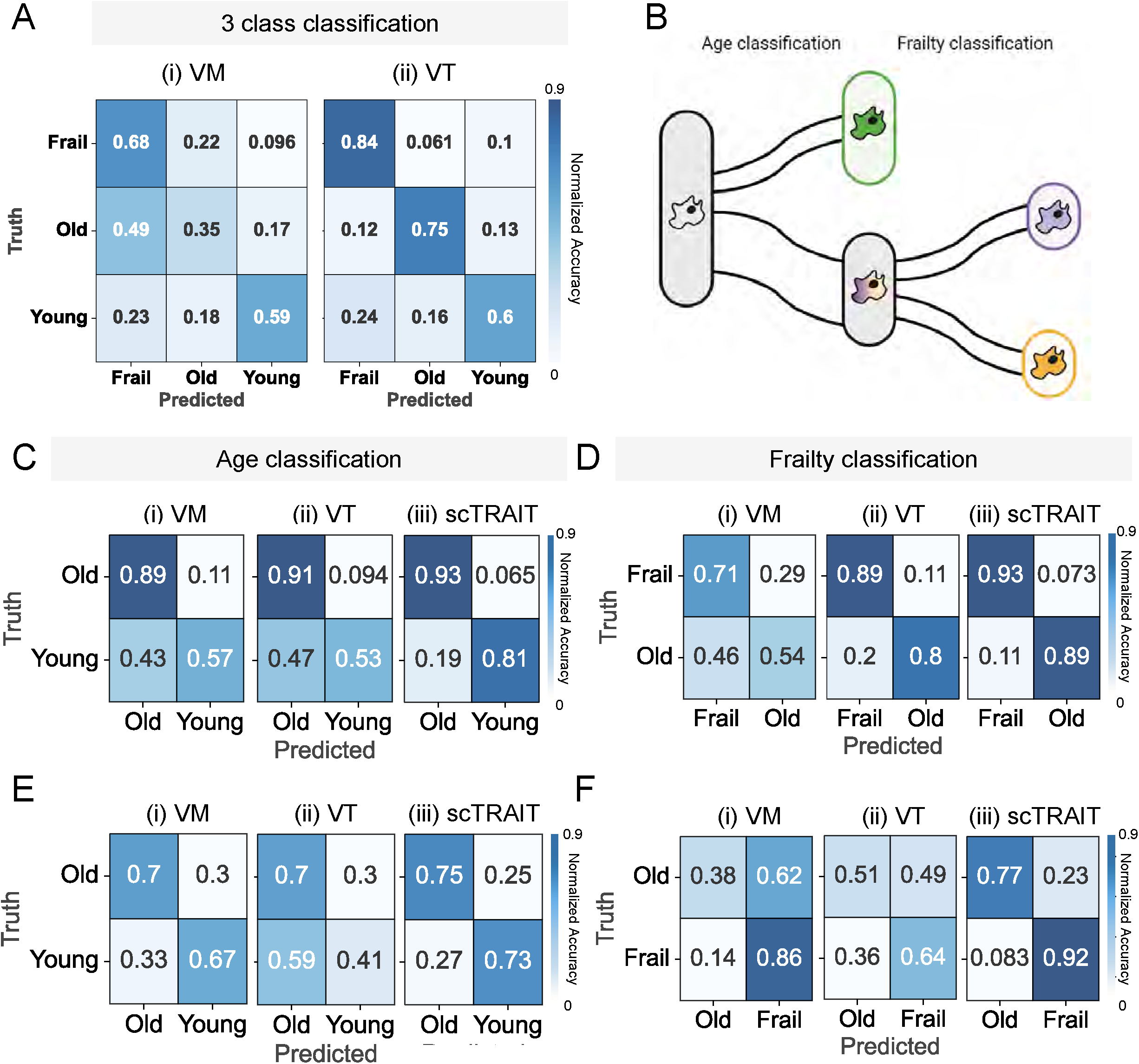
**A**. Normalized confusion matrices from two benchmark models vanilla morphodynamics (VM) and vanilla trajectory (VT) for 3 class classification. B. Schematic of two-step prediction strategy for predicting age and frailty status separately. First, we predict age based on young or old group (age classification), then predict frailty status within the old cohorts (frailty classification). **C-D**. Normalized confusion matrices from two benchmark models and scTRAIT for the age (C) and frailty (D) classification tasks. **E-F**. Normalized confusion matrices on the technical replicate dataset for age (E) and frailty (F) classification (scTRAIT AUC=0.82, 0.91 for age and frailty, respectively).

**Supplementary Figure 9.**
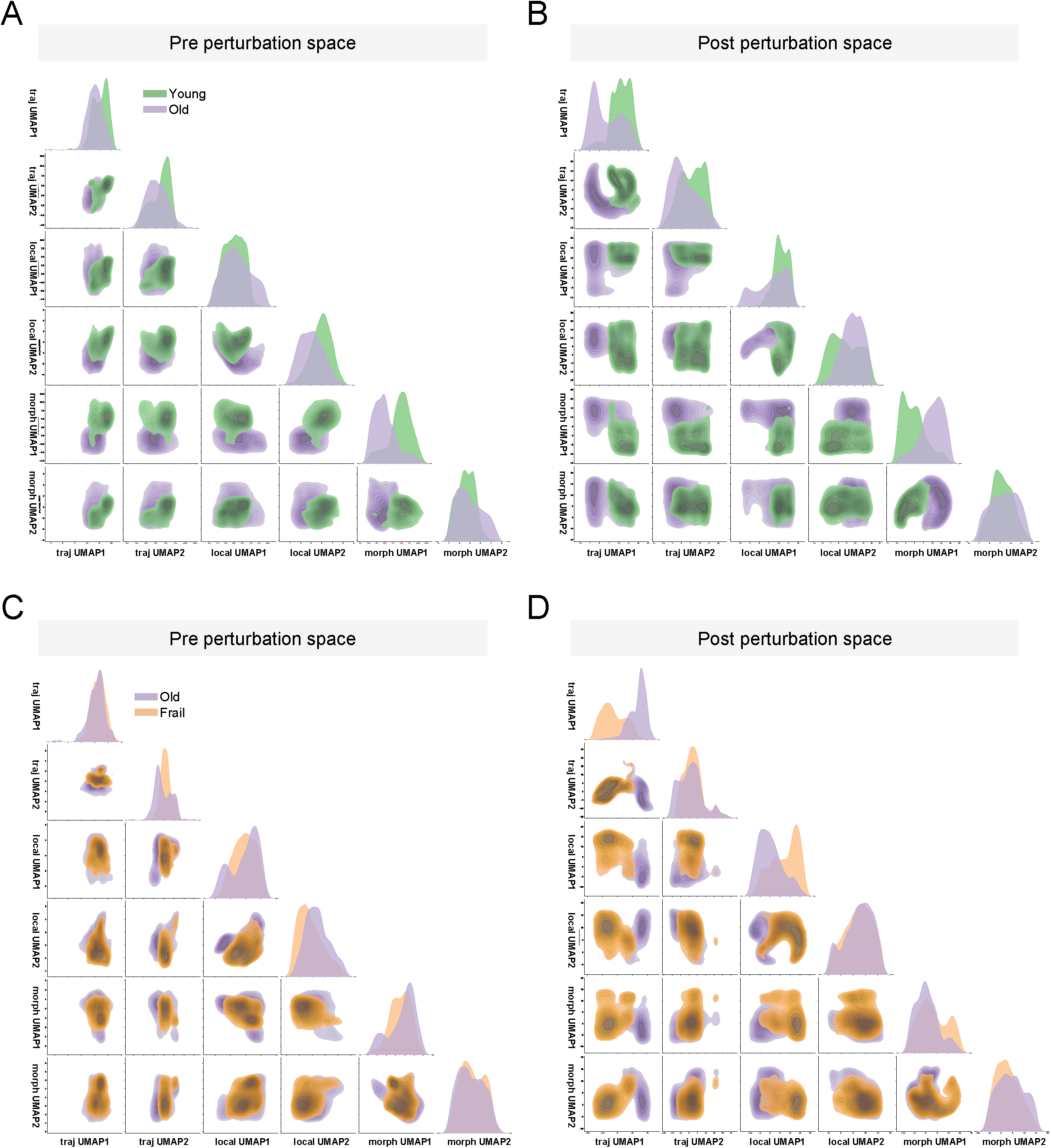
**A-B**. Pairwise two-dimensional UMAP projection space constructed by latent vectors from each motility (traj), colocalization (local) and morphodynamics (morpho) behavior modality based on pre perturbation (A) or post perturbation (B) outcome for age classification task. **C-D**. Corresponding pairwise UMAP space for frailty classification task exploring pre perturbation (C), or post perturbation (D) space.

**Supplementary Figure 10.**
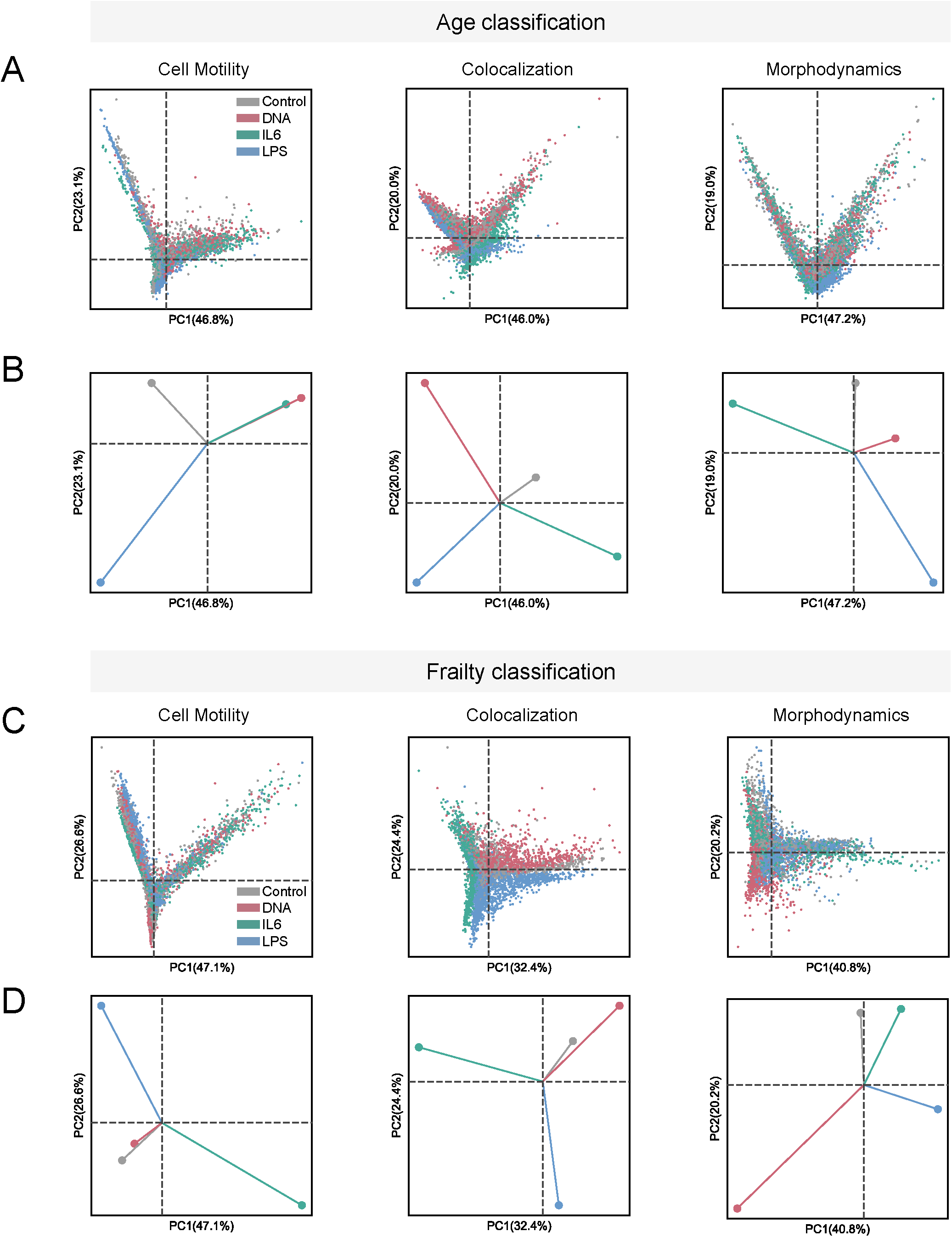
**A-B**. Single-cell (A), or global (B) perturbation embedding PCA space for each behavior modality in age classification task. Corresponding **C-D**. single-cell (C) or global (D) perturbation embedding PCA space in frailty classification task. Global embedding is calculated by taking the centroid of PCA coordinates for each perturbation.

**Supplementary Figure 11.**
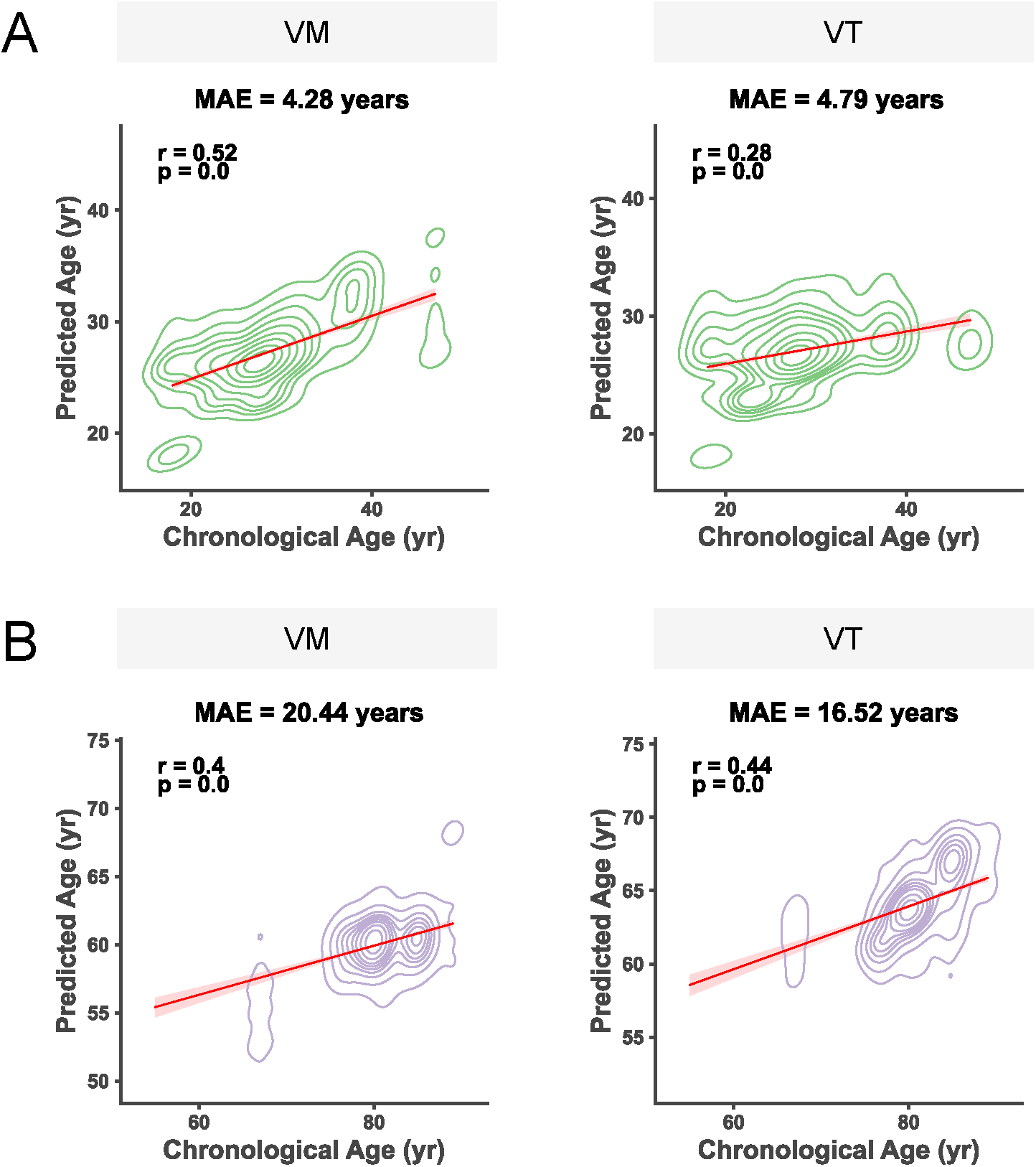
**A-B**. Predicted age distribution over chronological age from two benchmark models on young (A), and old (B) cells. Corresponding MAE and Pearson’s correlation coefficient r is shown.

**Supplementary Figure 12.**
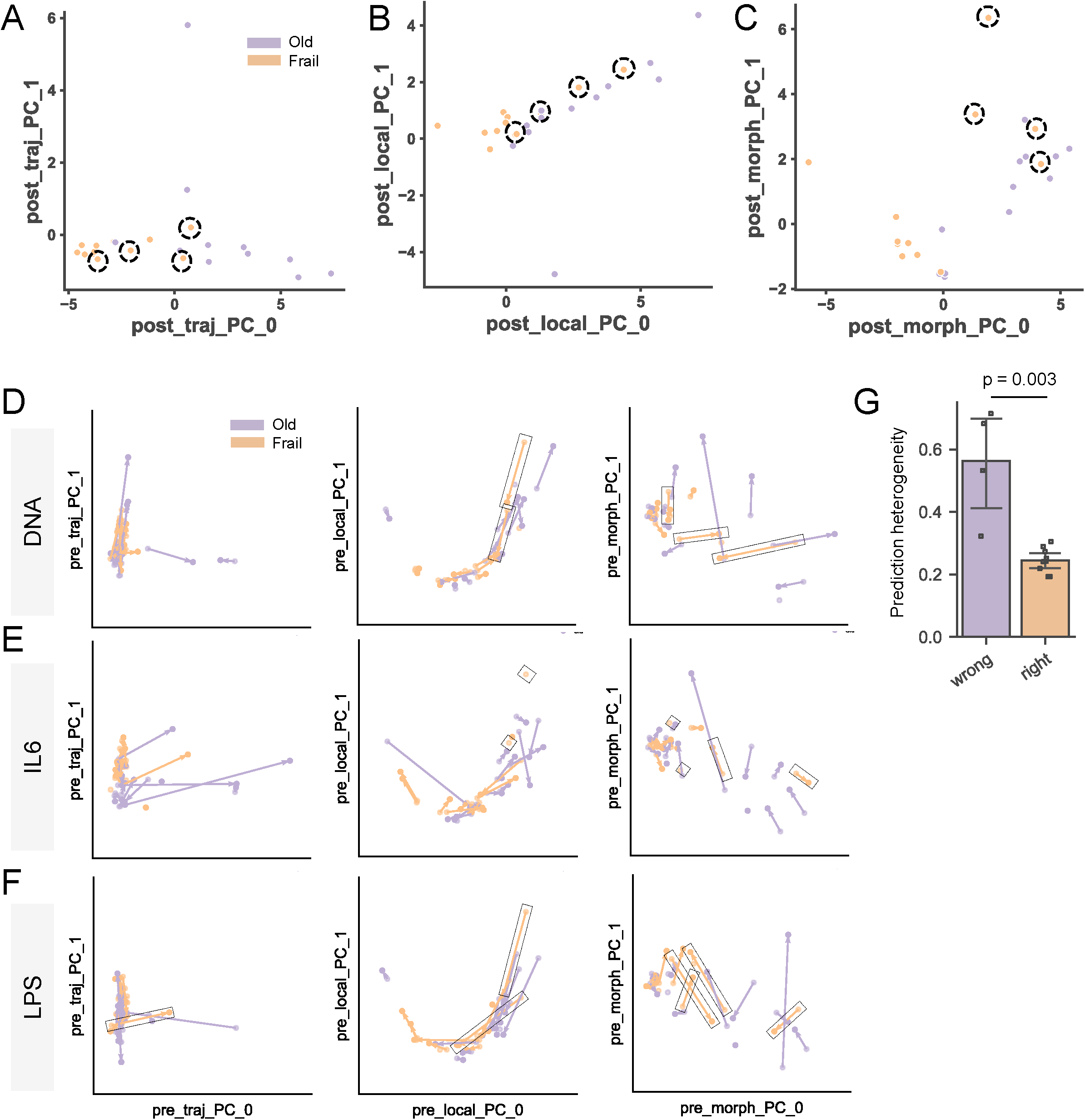
**A-C**. Projection of donors onto two-dimensional PCA space of post perturbation behavioral states defined by motility (A), colocalization (B) and morphodynamics (C). Dotted circle represents the frail donors that are predicted as non-frail old based on scTRAIT. **D-F**. Two-dimensional PCA plot showing the responses of non-frail old and frail cohorts on each perturbation DNA (D), IL6 (E) and LPS (F) based on behavior modalities from left to right motility, colocalization and morphodynamics. Vectors depict each donor’s perturbation response on the behavior space. Dotted rectangle indicates frail donor s that are predicted as non-frail old based on scTRAIT. **G**. Predicted heterogeneity calculated by coefficient of variance of predicted age within frail donors that scTRAIT predicted non-frail (wrong) or frail (right) (wrong N = 4, right N = 9 donors).

**Supplementary Figure 13.**
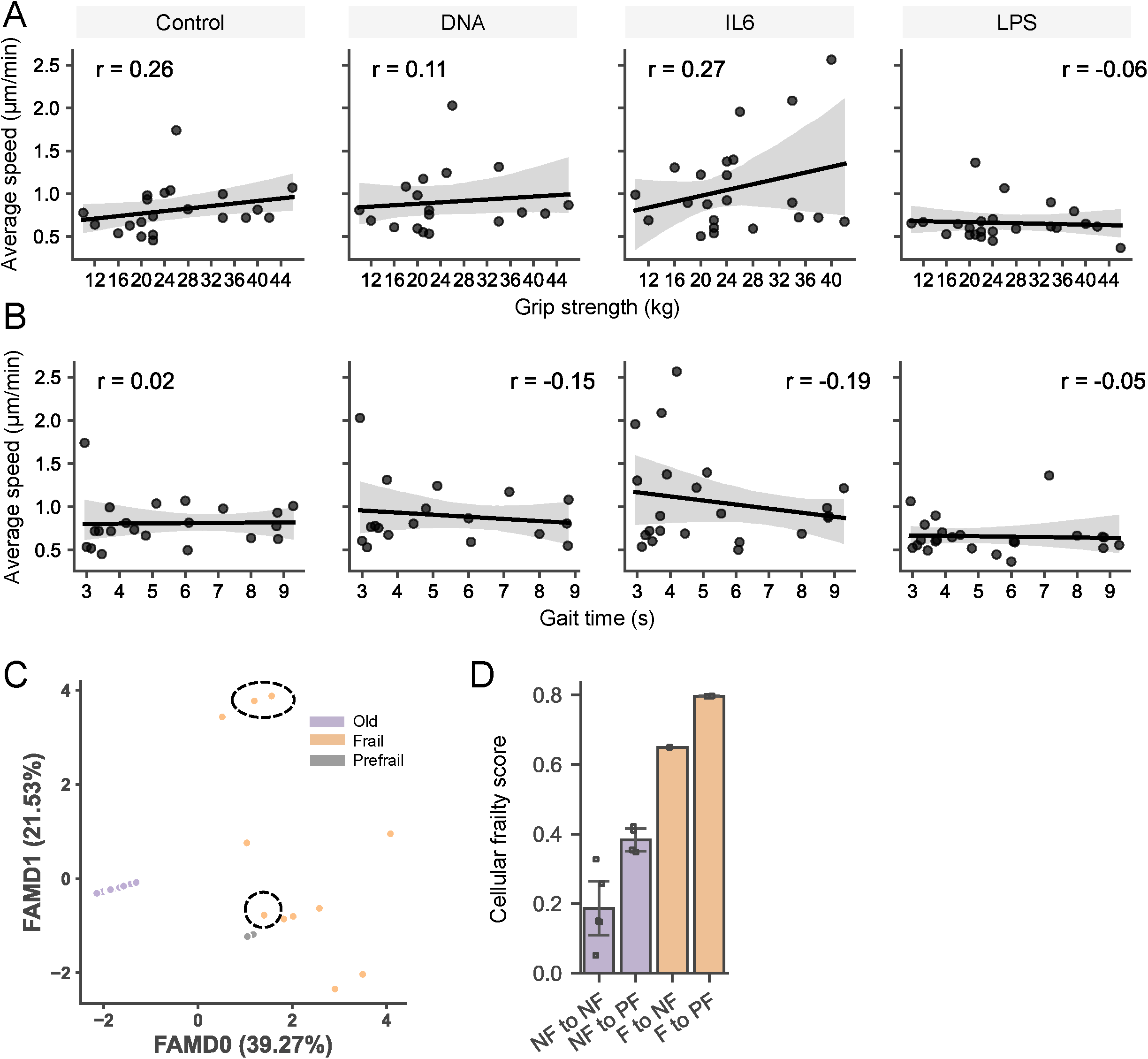
**A-B**. Regression plot of **A**. grip strength and **B**. gait time over average speed across control and 3 perturbation conditions. **C**. Two-dimensional factor analysis of mixed data (FAMD) of clinical measurements of donors from non-frail old, prefrail and frail group. Dotted circle represents the frail donors that are predicted as non-frail old based on scTRAIT. **D**. Cellular frailty score of donors transitioned from NF (Non-frail) to NF, NF to PF (Pre-frail), F (Frail) to NF and F to PF.

