## Supplementary Figures for "Monocytes are biological sensors of aging and frailty in humans"

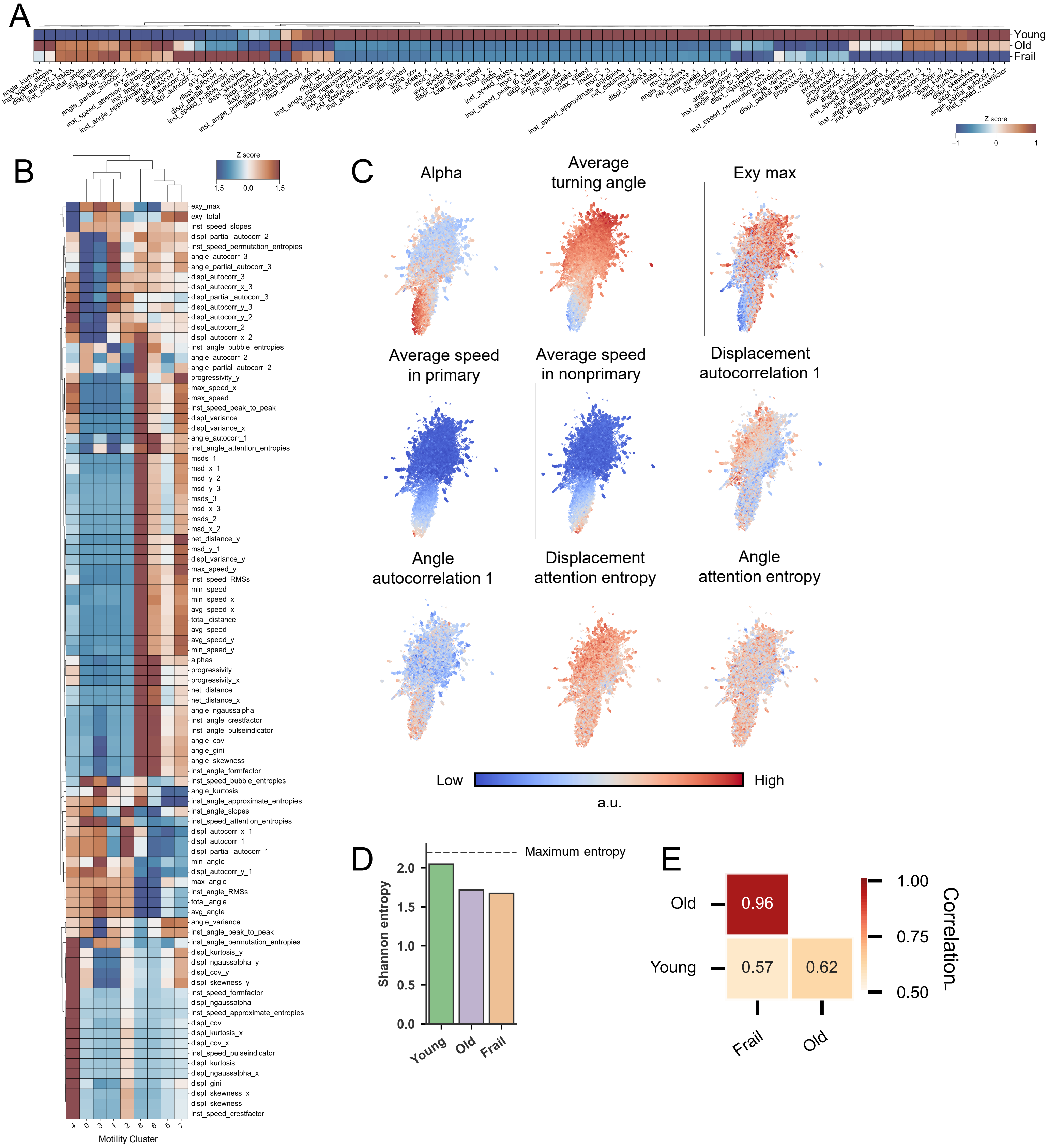


**Supplementary Figure 1. A.** Z scores across motility parameters for each age group. Hierarchical clustering is based on the Euclidean distance using Ward method groups similar Z scores of features. **B.** Z scores across motility parameters for each motility cluster. Hierarchical clustering is based on the Euclidean distance using Ward method groups similar Z scores of features. **C.** Projection of various motility features onto UMAP space. **D.** Shannon entropy to quantify heterogeneity of motility states. The dotted line indicates the maximum entropy that can be calculated from nine motility cluster distributions. **E.** Cross correlation based on the fraction of motility clusters.

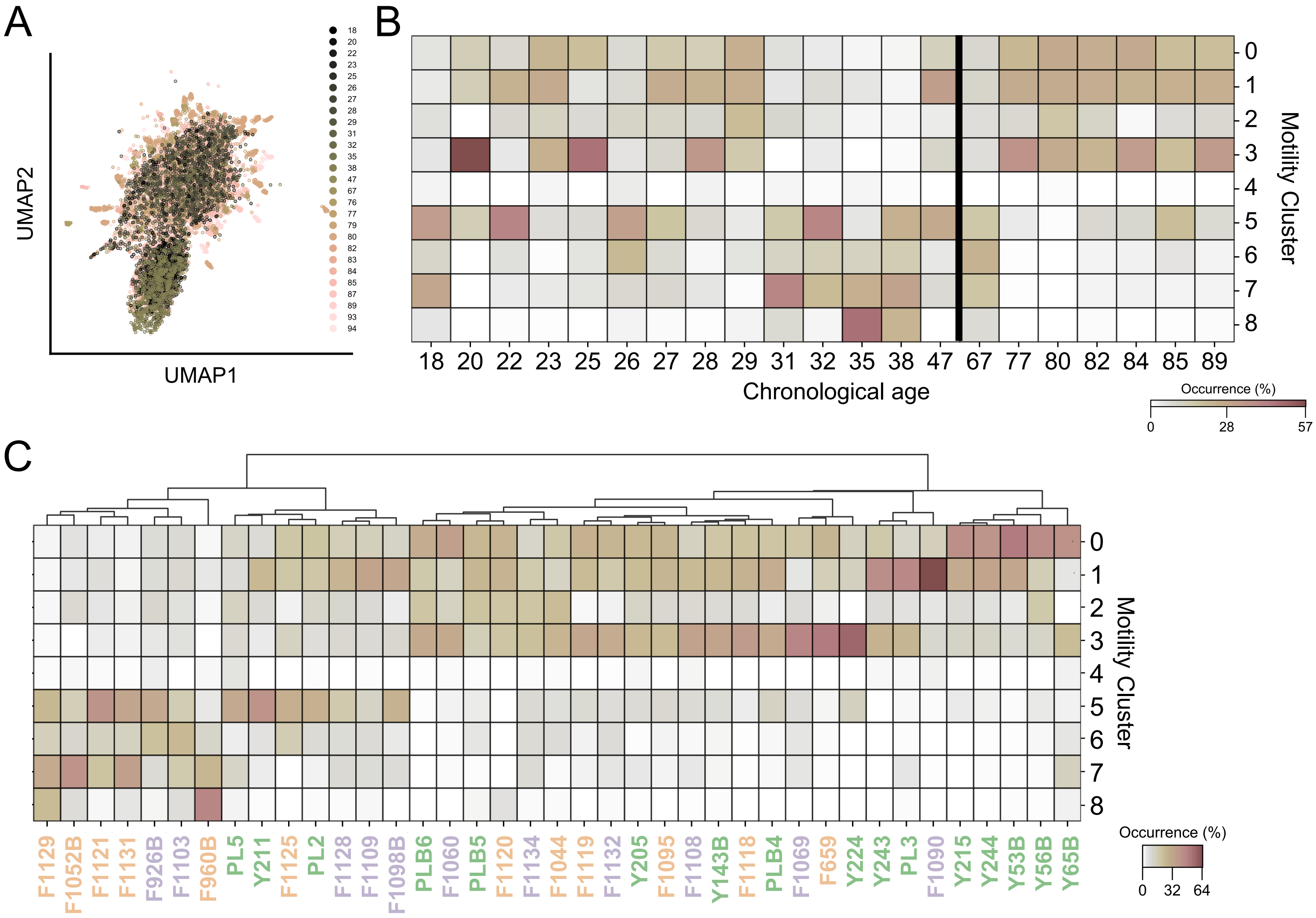


**Supplementary Figure 2. A.** Spectrum of age projected onto the UMAP space describing the donor-to-donor variation within the young, old and frail groups for control condition. **B.** Motility cluster distribution for each age under control, suggesting non-linear and heterogeneous cluster enrichment patterns especially within the young. A bold line distinguishes the young and old group. **C.** MC distribution across all donors in control condition highlights the donor-to-donor variation in the motility patterns within each age group. Green, purple, orange colors represent young, old, frail groups respectively.

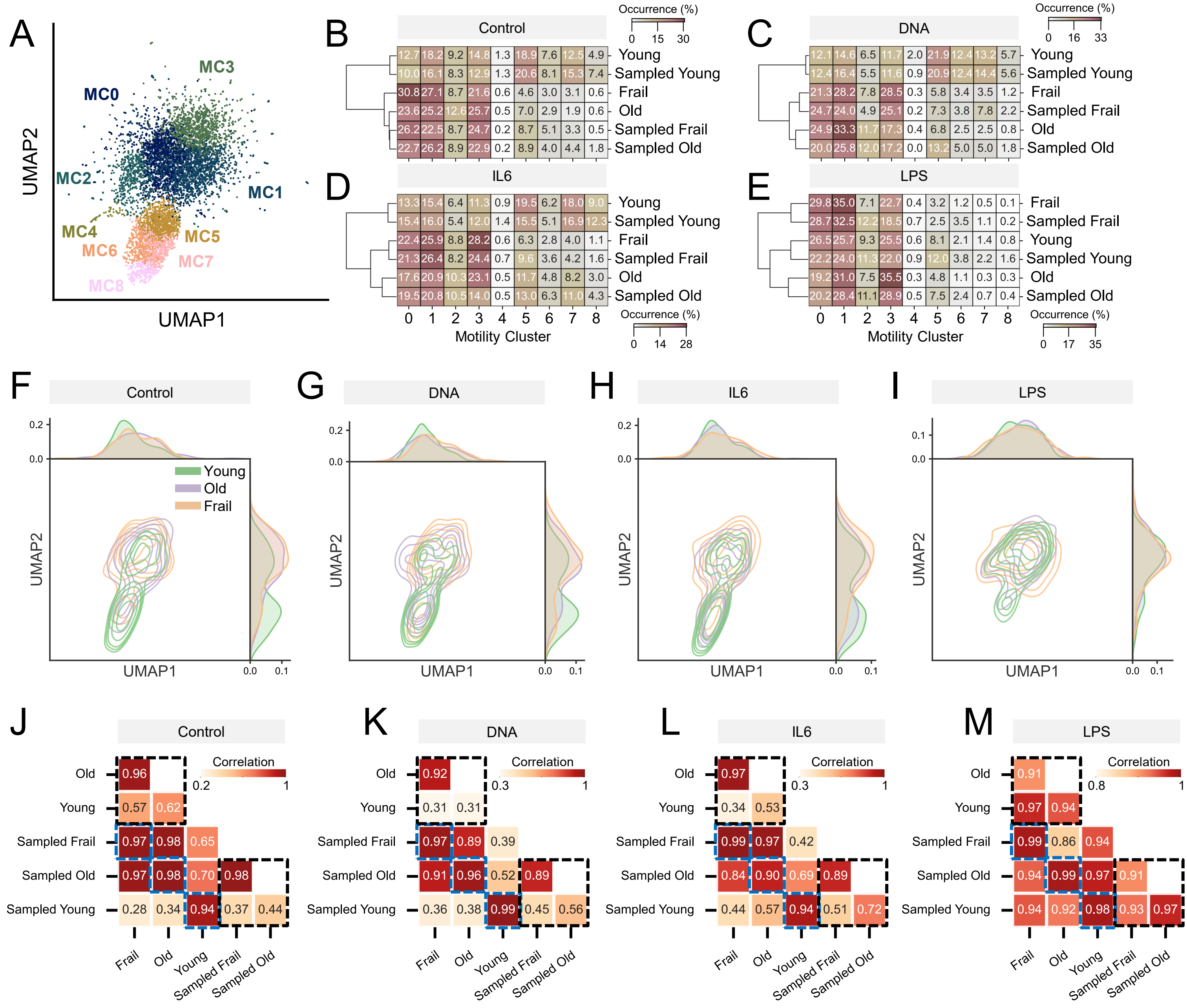

**Supplementary Figure 3. A.** 2D UMAP representation of nine motility clusters in the sampled space (sampling n = 50 each donor). **B-E.** Heatmap showing the fractional occurrence of nine MCs across the age groups and sampled age groups exposed to control (B), DNA (C), IL6 (D), and LPS (E). Hierarchical clustering is based on the Euclidean distance using the Ward method. The sum of rows equal 1. **F-I.** 2D UMAP KDE representation of sampled space across age groups exposed to control (F), DNA (G), IL6 (H), and LPS (I) conditions. 1D distributions of single cells for each at the top and right margins. **J-M.** Cross correlation based on the fraction of motility clusters exposed to control (J), DNA (K), IL6 (L), and LPS (M). Top left and bottom right dotted black box represent comparison within original and sampled space, respectively. Blue dotted box represents comparison between original and sampled space for each age group.

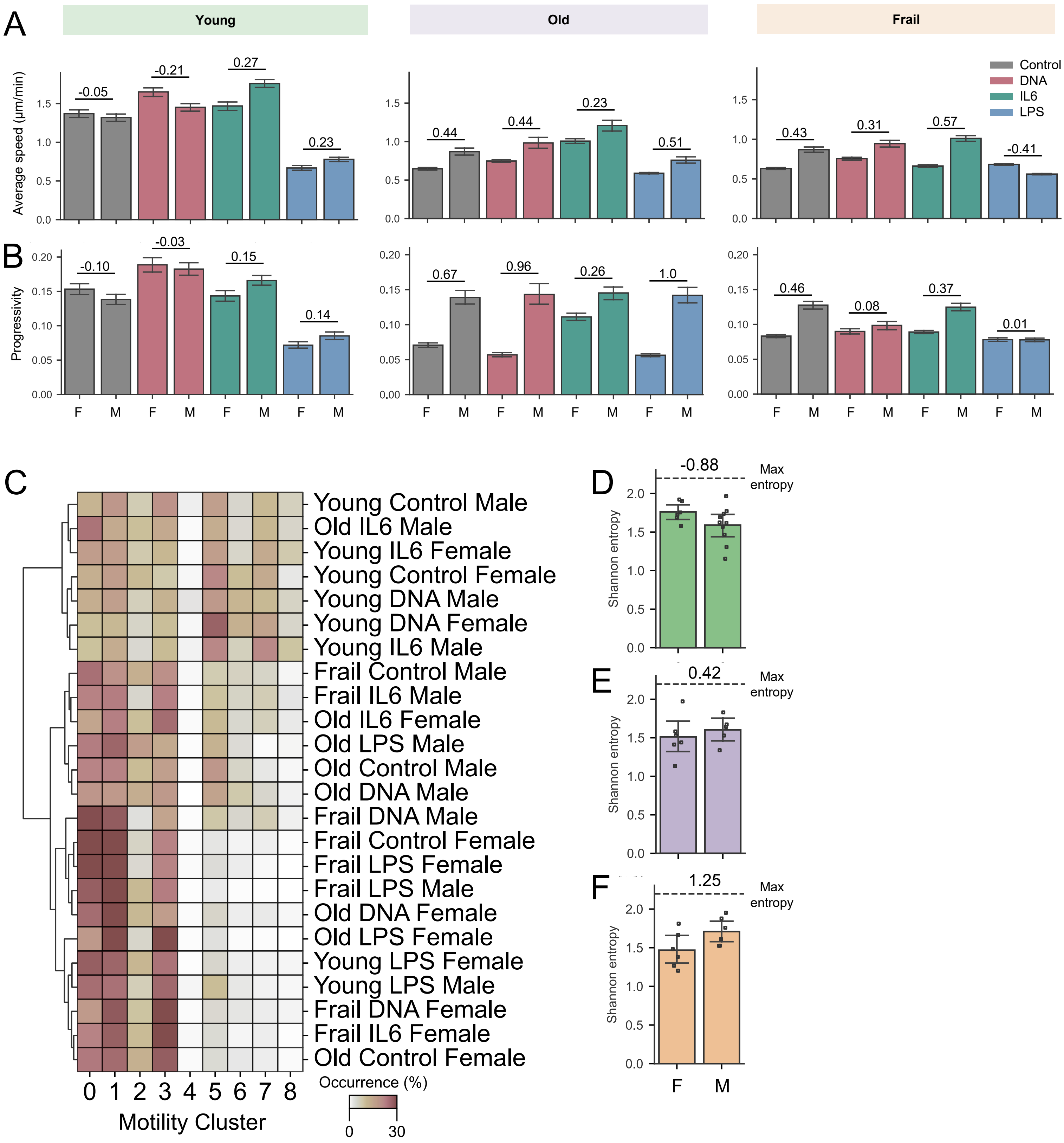


**Supplementary Figure 4. A-B.** Bar plot comparing female (F) and male (M) with respect to control, DNA, IL6, and LPS based on average speed (A), and progressivity (B). Annotated values represent Cohen's d. **C.** MC distribution of cells that are categorized combining age groups, perturbation and gender. Hierarchical clustering is based on the Euclidean distance using Ward method. **D-F.** Shannon entropy in control group comparing female (F) and male (M) within young (D), old (E) and frail (F) group (mean ± 95% C.I.). Annotated values represent Cohen's d (young female N=6, young male N=10, old female N=6, old male N=5, frail female N=6, frail male N=6 donors).

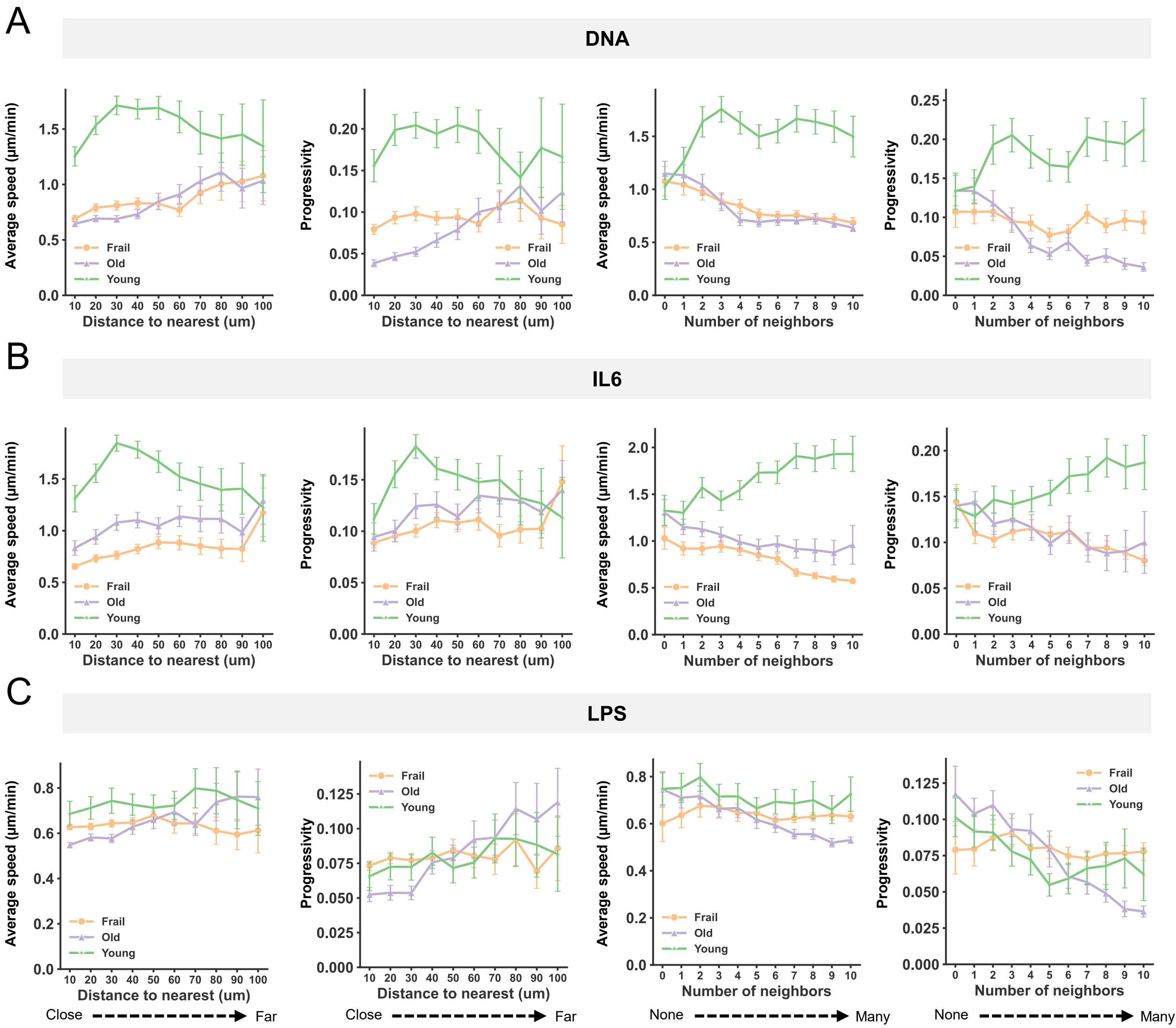


**Supplementary Figure 5. A-C.** Average speed and progressivity over the distance to the average nearest cell with 10 μm bin (mean ± 95% C.I.) and over the average number of neighbors within the 100 μm range with 1 cell bin (mean ± 95% C.I.) under DNA (A), IL6 (B), LPS (C) conditions.

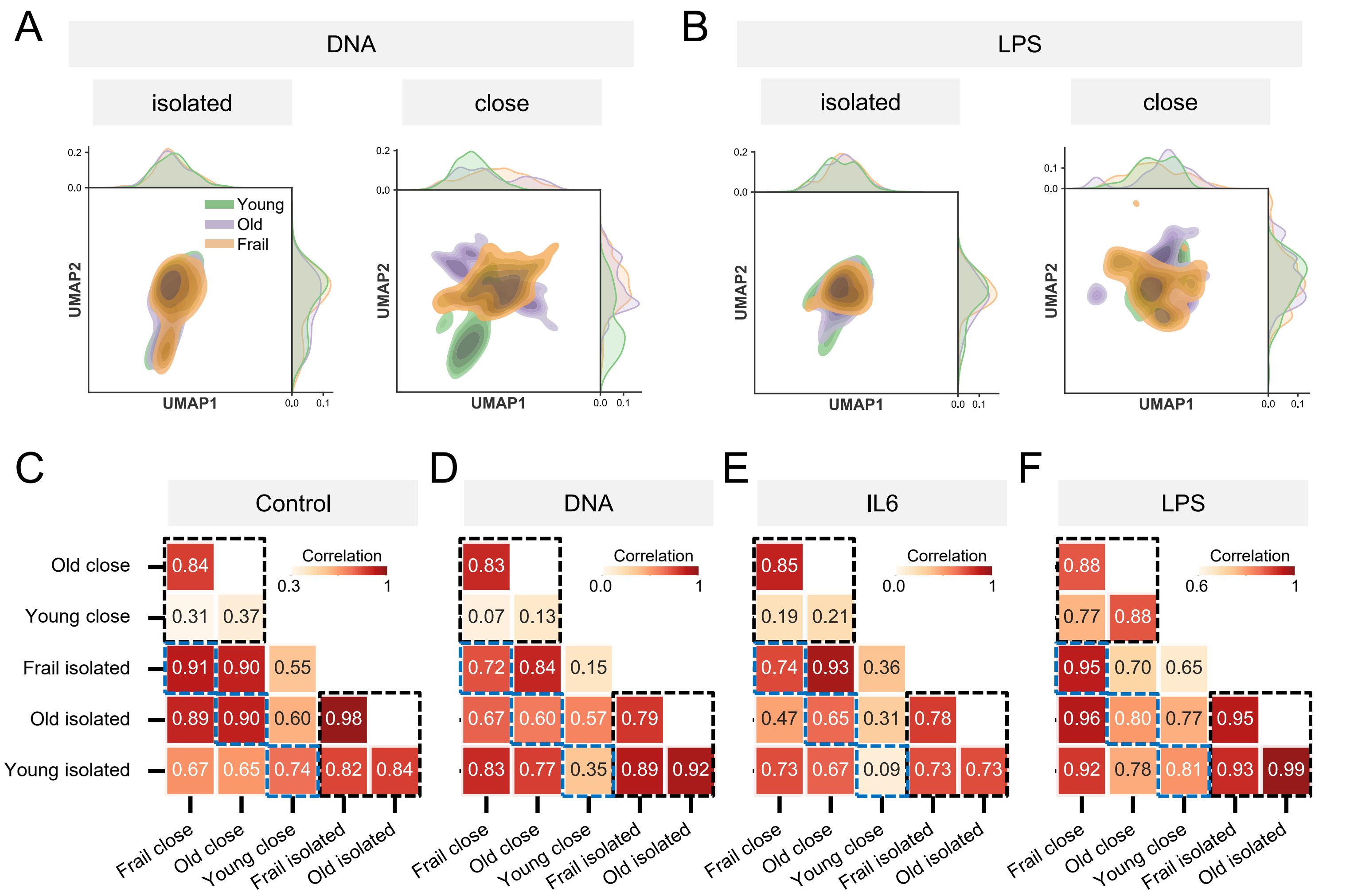


**Supplementary Figure 6. A-B.** UMAP projection of motility distribution comparing isolated and close conditions within DNA (A) and LPS (B) treated experiments. **C-F.** Cross correlation based on the fraction of motility clusters for 3 age groups with each isolated and close condition within Control (C), DNA (D), IL6 (E), and LPS (F) treated samples. Top left and bottom right dotted black box represent comparison within close and isolated monocytes, respectively. Blue dotted box represents comparison between close and isolated cells for each age group.

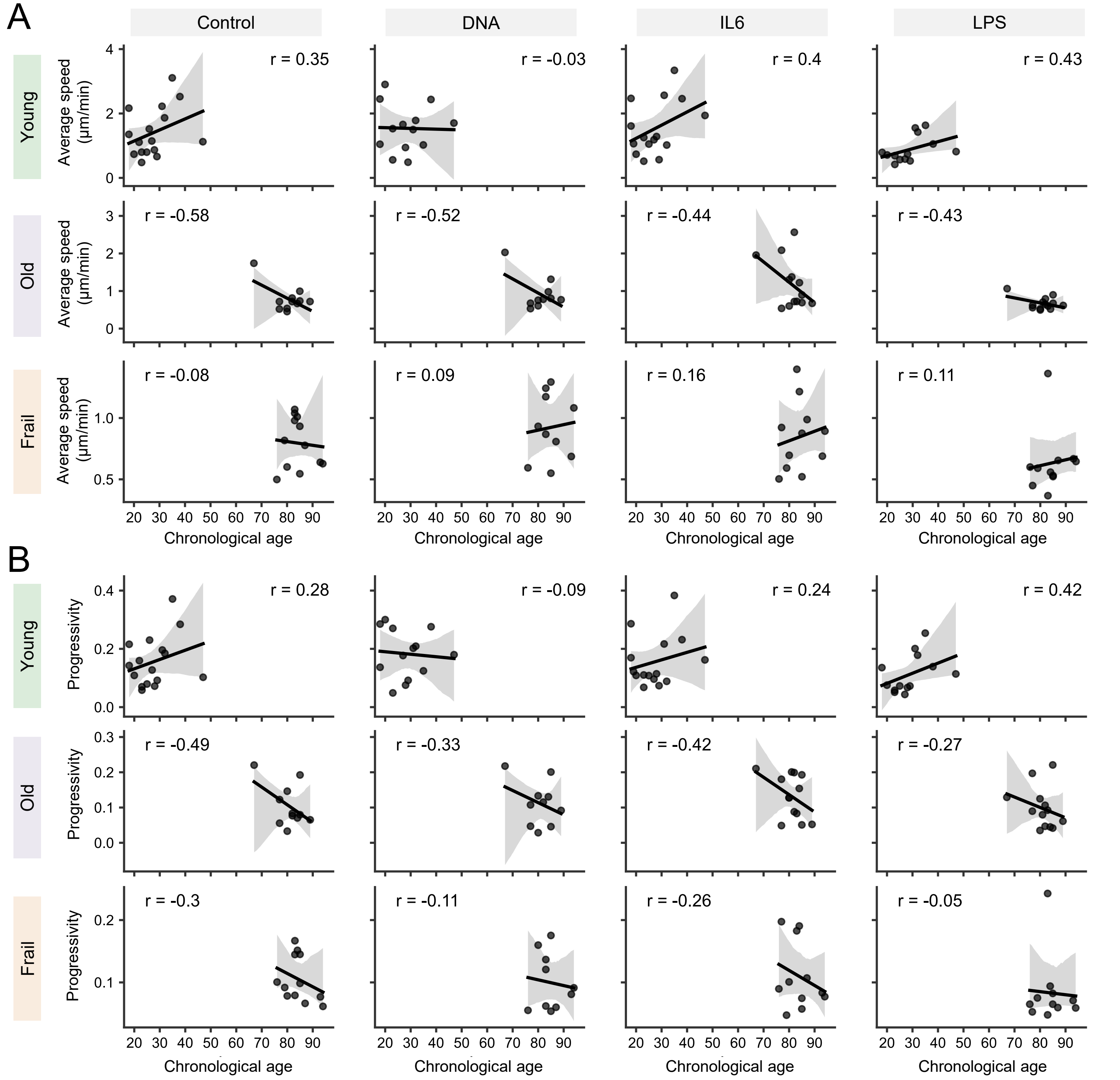


**Supplementary Figure 7. A-B.** Regression plot of average speed (A) and progressivity (B) over chronological age across 3 age groups. Annotated r refers to Pearson's correlation coefficient.

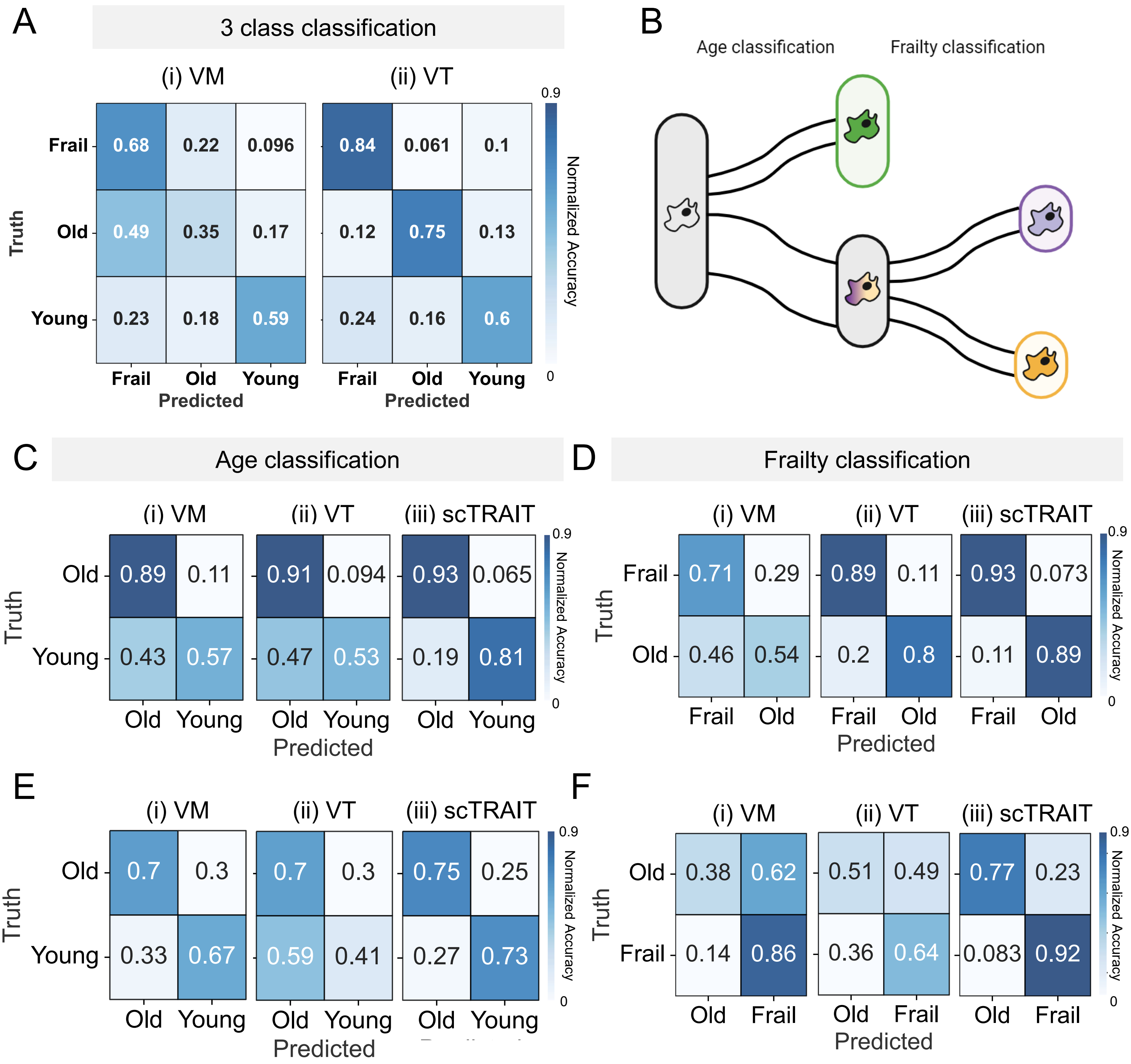


**Supplementary Figure 8. A.** Normalized confusion matrices from two benchmark models vanilla morphodynamics (VM) and vanilla trajectory (VT) for 3 class classification. **B.** Schematic of two-step prediction strategy for predicting age and frailty status separately. First, we predict age based on young or old group (age classification), then predict frailty status within the old cohorts (frailty classification). **C-D.** Normalized confusion matrices from two benchmark models and scTRAIT for the age (C) and frailty (D) classification tasks. **E-F.** Normalized confusion matrices on the technical replicate dataset for age (E) and frailty (F) classification (scTRAIT AUC=0.82, 0.91 for age and frailty, respectively).

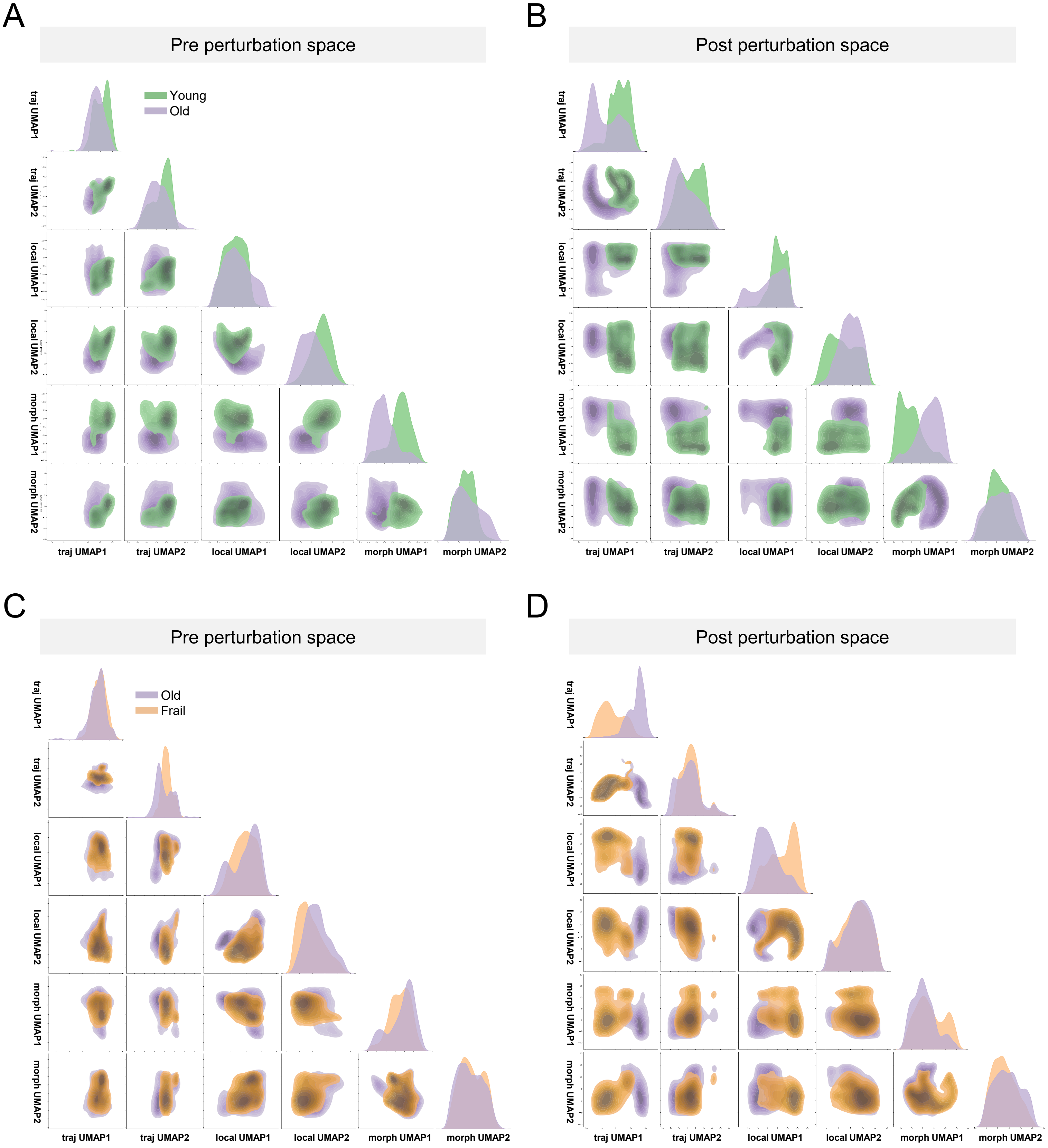


**Supplementary Figure 9. A-B.** Pairwise two-dimensional UMAP projection space constructed by latent vectors from each motility (traj), colocalization (local) and morphodynamics (morpho) behavior modality based on pre perturbation (A) or post perturbation (B) outcome for age classification task. **C-D.** Corresponding pairwise UMAP space for frailty classification task exploring pre perturbation (C), or post perturbation (D) space.

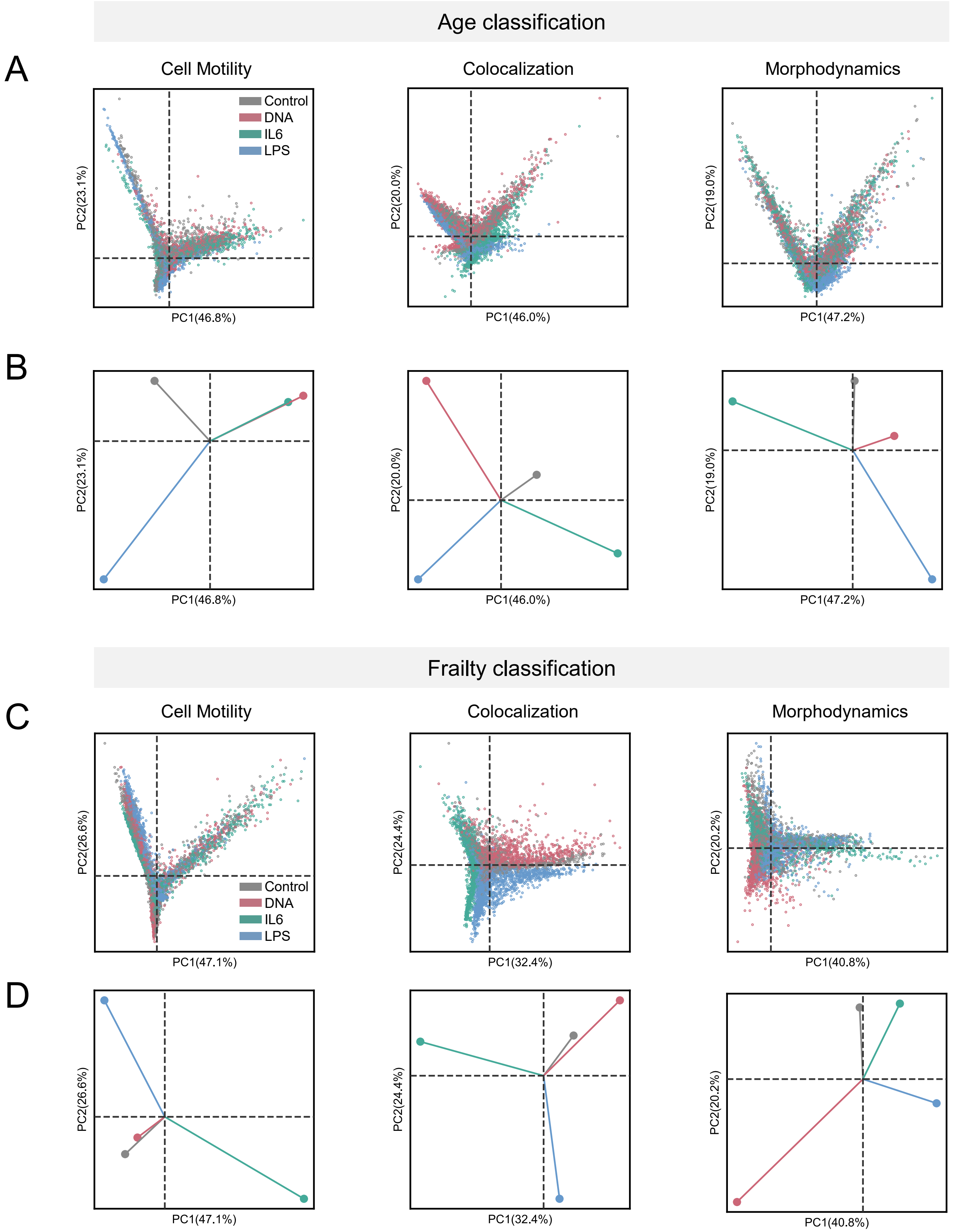


**Supplementary Figure 10. A-B.** Single-cell (A), or global (B) perturbation embedding PCA space for each behavior modality in age classification task. Corresponding **C-D.** single-cell (C) or global (D) perturbation embedding PCA space in frailty classification task. Global embedding is calculated by taking the centroid of PCA coordinates for each perturbation.

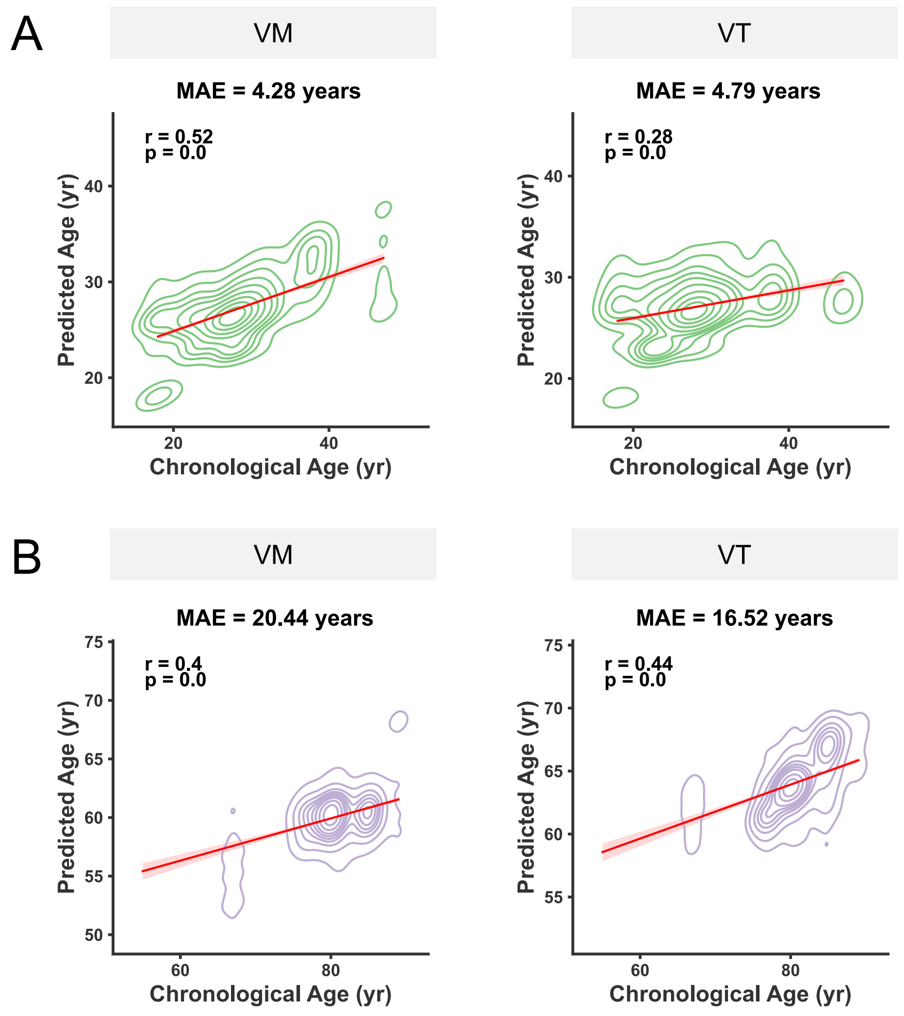


**Supplementary Figure 11. A-B.** Predicted age distribution over chronological age from two benchmark models on young (A), and old (B) cells. Corresponding MAE and Pearson's correlation coefficient r is shown.

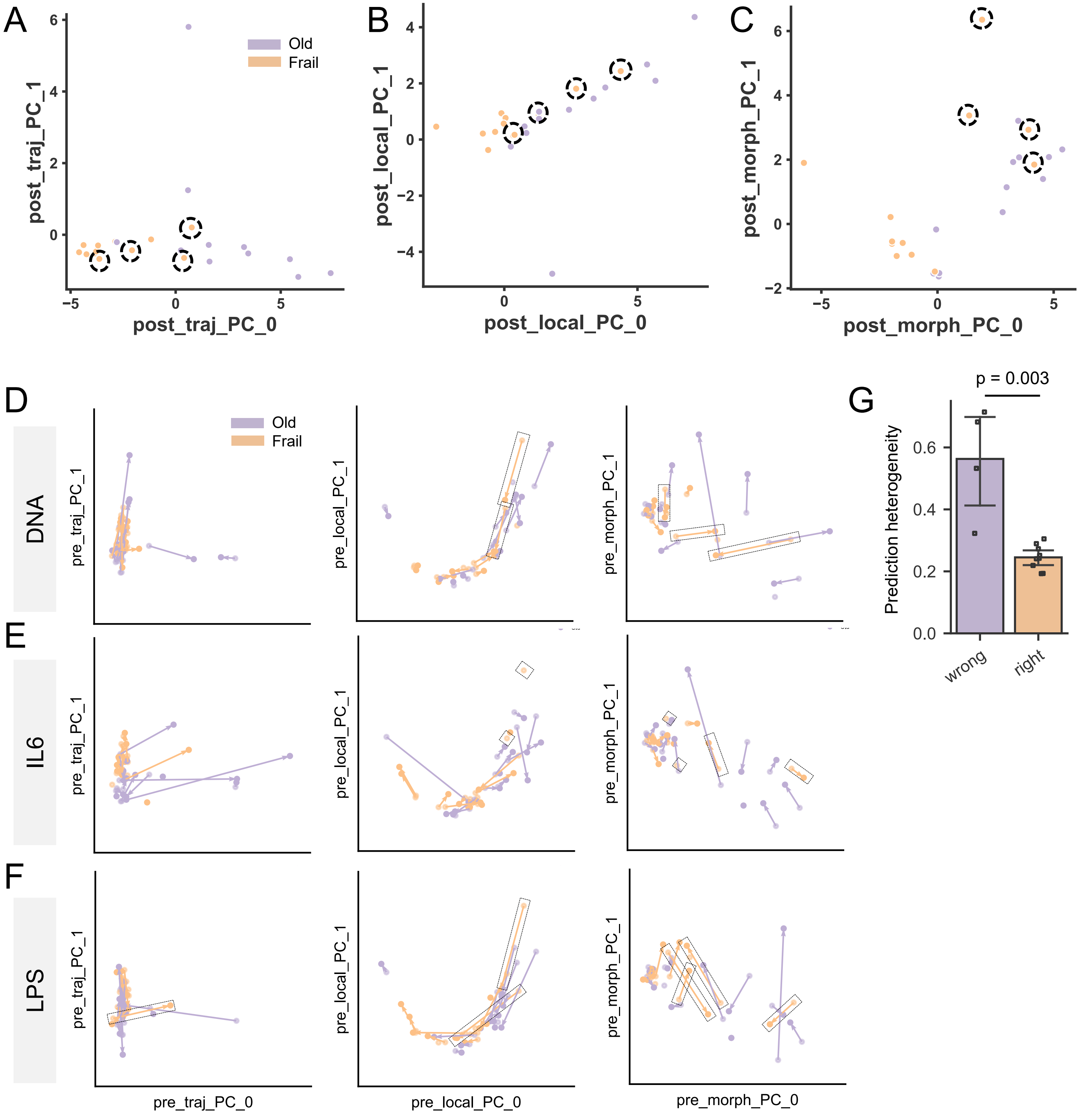


**Supplementary Figure 12. A-C.** Projection of donors onto two-dimensional PCA space of post perturbation behavioral states defined by motility (A), colocalization (B) and morphodynamics (C). Dotted circle represents the frail donors that are predicted as non-frail old based on scTRAIT. **D-F.** Two-dimensional PCA plot showing the responses of non-frail old and frail cohorts on each perturbation DNA (D), IL6 (E) and LPS (F) based on behavior modalities from left to right motility, colocalization and morphodynamics. Vectors depict each donor's perturbation response on the behavior space. Dotted rectangle indicates frail donor s that are predicted as non-frail old based on scTRAIT. **G.** Predicted heterogeneity calculated by coefficient of variance of predicted age within frail donors that scTRAIT predicted non-frail (wrong) or frail (right) (wrong N = 4, right N = 9 donors).

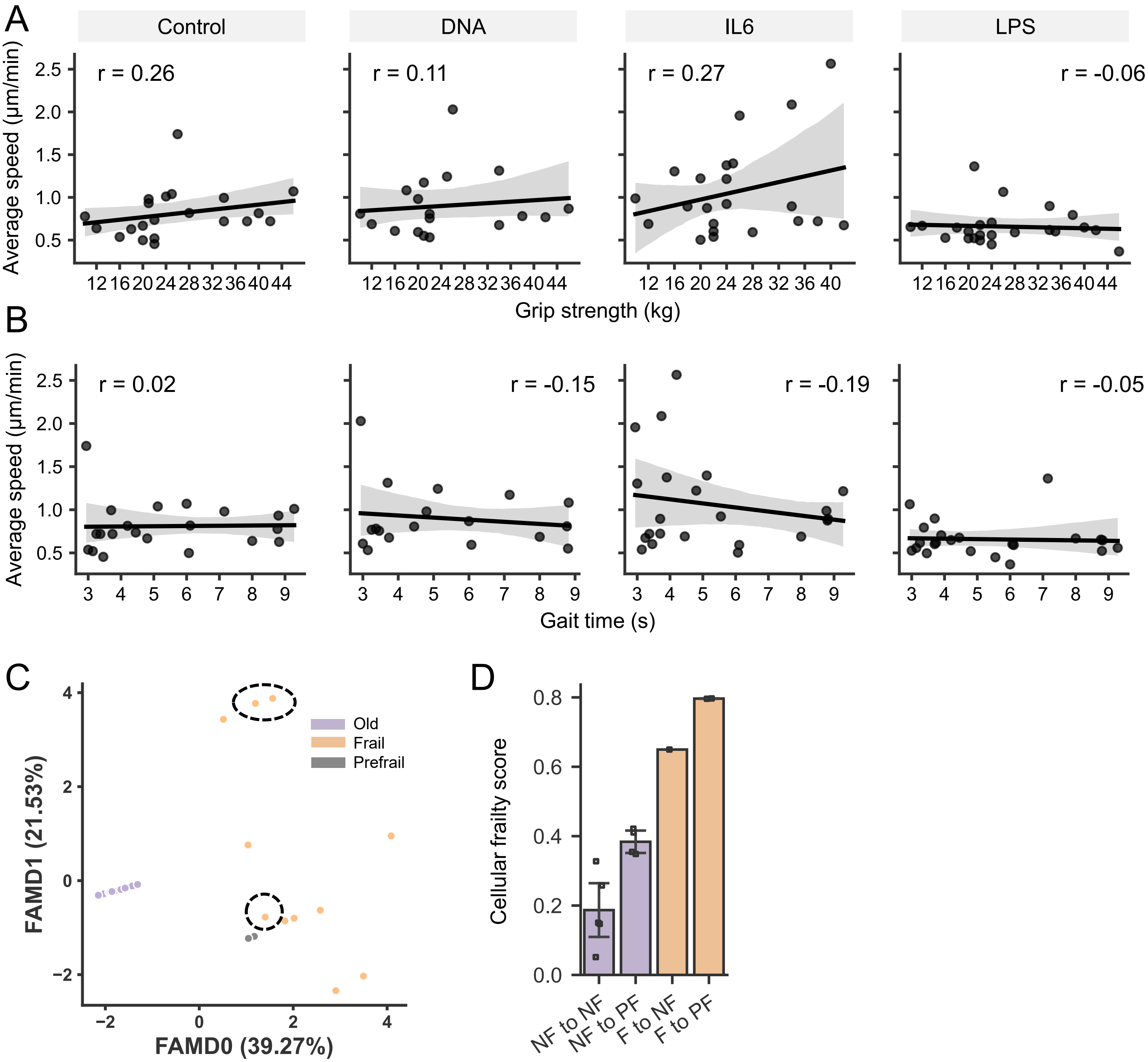


**Supplementary Figure 13. A-B.** Regression plot of **A.** grip strength and **B.** gait time over average speed across control and 3 perturbation conditions. **C.** Two-dimensional factor analysis of mixed data (FAMD) of clinical measurements of donors from non-frail old, prefrail and frail group. Dotted circle represents the frail donors that are predicted as non-frail old based on scTRAIT. **D.** Cellular frailty score of donors transitioned from NF (Non-frail) to NF, NF to PF (Pre-frail), F (Frail) to NF and F to PF.
